# A systems model of alternating theta sweeps via firing rate adaptation

**DOI:** 10.1101/2024.09.13.612841

**Authors:** Zilong Ji, Tianhao Chu, Si Wu, Neil Burgess

**Author notes:** These authors contribute equally to this work (Zilong Ji, Tianhao Chu).

## Abstract

Place and grid cells provide a neural system for self-location and tend to fire in sequences within each cycle of the hippocampal theta rhythm when rodents run on a linear track. These sequences correspond to the decoded location of the animal sweeping forward from its current location (“theta sweeps”). However recent findings in open-field environments show alternating left-right theta sweeps, and propose a circuit for their generation. Here, we present a computational model of this circuit, comprising head direction cells, conjunctive grid x direction cells, and pure grid cells, based on continuous attractor dynamics, firing rate adaptation, and modulated by the medial-septal theta rhythm. Due to firing rate adaptation, the head-direction ring attractor exhibits left-right sweeps coding for internal direction, providing an input to the grid cell attractor network shifted along the internal direction, via an intermediate layer of conjunctive grid x direction cells, producing left-right sweeps of position by grid cells. Our model explains the empirical findings, including the alignment of internal position and direction sweeps and the dependence of sweep length on grid spacing. It makes predictions for theta-modulated head-direction cells, including specific relationships between theta phase precession during turning, theta skipping, anticipatory firing and directional tuning width. These predictions are verified in experimental data from anteroventral thalamus. The model also makes several predictions for the relationships between position and direction sweeps, running speed and dorsal-ventral location within the entorhinal cortex. Overall, a simple intrinsic mechanism explains the complex theta dynamics of the spatial circuit, with testable predictions.

## Introduction

The entorhinal-hippocampal system plays a crucial role in spatial navigation and goal-directed planning [1, 2, 3]. Various spatially tuned cells, including head direction [4], place [5], grid [6] and boundary vector cells [7] support the underlying computational processes involved in these functions. Beyond spatial coding, place and grid cells exhibit a temporal coding feature, i.e., theta phase precession, where the sequential firing of individual neurons occurs progressively earlier in relation to the phase of the ongoing theta rhythm driven by inputs from the medial septum [8, 9]. At the populational level, phase precessing cells underlie forward-directed theta sweeps in linear track environments [10], alternating sweeps in T-maze environments [11, 12] and left-right-alternating sweeps in open-field environments [13].

While many network models have been proposed to explain the generation of theta sweeps [14, 15, 16], they primarily model forward-directed sweeps along the running direction of the animal in linear track environments. This is because the cells are arranged in a one-dimensional manner to fit the environmental setting. It remains unclear how these models can be extended to account for theta sweeps in less constrained environments, such as open fields (but see [17] for theta sweeps in T-mazes). Inspired by a recent study demonstrating left-right-alternating theta sweeps from recordings of medial entorhinal cortex (MEC) grid cells in open-field environments[13], we developed a continuous attractor network model to illustrate theta sweeps in such environments. Our model comprises a three-layer network of head direction cells, conjunctive grid x head direction cells (hereafter referred to as conj-grid cells) and pure grid cells (Fig. 1a). The head direction cells form a ring attractor (hereafter referred to as the HD attractor), where the activity bump represents the animal’s internal direction. This HD attractor activates downstream conj-grid cells whose preferred directional tuning aligns with the internal direction. The conj-grid cells then carry position-dependent input with an offset along the internal direction onto the pure grid cells attractor network (hereafter referred to as the GC attractor), driving the activity bump in the GC attractor to move. Due to firing rate adaptation and medial-septal theta modulation, the internal direction represented by the activity bump in the HD attractor sweeps from side to side of the animal’s head axis at theta rhythm. This consequently results in left-right-alternating bump sweeps in the GC attractor via the conj-grid cells.

**Figure 1:**
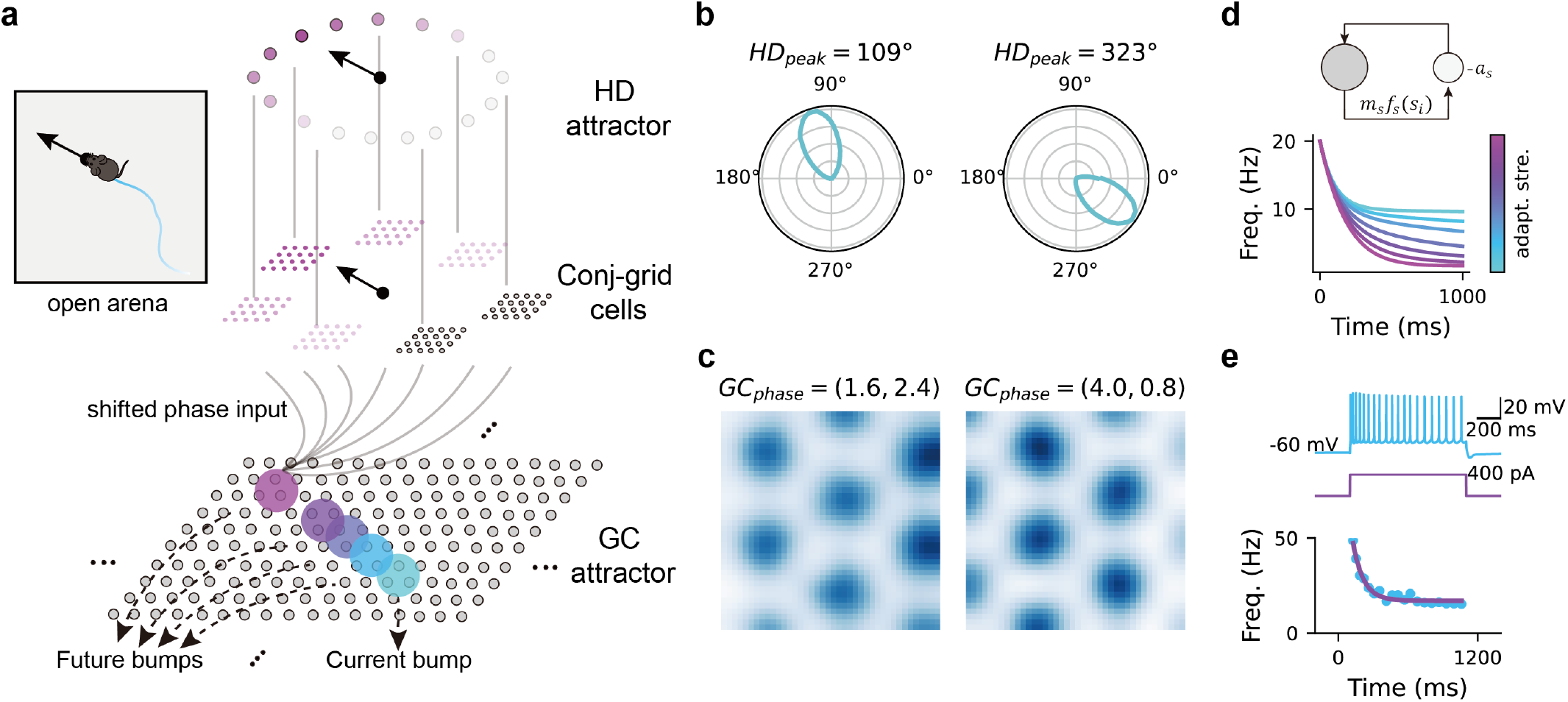
The continuous attractor network model with firing rate adaptation. **a**. illustration of the computation model, consisting of a head direction ring attractor (HD attractor) and a grid cell 2-dimension attractor (GC attractor). The HD attractor receives input of the animal’s head direction, projecting directional input to the conjunctive head × grid cells, which further send a shifted phase input along the direction of head axis to drive the activity bump in the GC attractor. **b**. two example head direction cells in the HD attractor. **c**. two example grid cells in the GC attractor. **d**. top: illustration of the firing rate adaptation which is a slow negative feedback modulation to cell firing. Bottom: cell firing frequency as a function of time when applying both a constant input current. Different colors represent different adaptation strength. **e**. train (blue) of action potentials elicited in a in-vitro whole-cell patch-clamp MEC cell during 1-second pulse injection (purple). Instantaneous firing frequency (blue dots) were shown at the bottom with an exponential fit (purple curves). The plot is adapted from Yoshida et al. [27] with permission.

This model explains many of the empirical findings reported in Vollan et al. [13], including: 1) the alignment of internal direction and internal location sweeps in separate attractor networks; 2) the linear relationship between sweep length and grid spacing; and 3) the increase of the alternation score with running speed. Moreover, with this model, we make several new predictions. These include how the modulatory theta input affects theta sweeps in both the head direction and the grid cell networks, how the head direction network controls theta sweeps in the downstream grid cell network, how animals’ behaviors (e.g., running speed; straight runs and turns) modulate theta sweep features, and how theta sweeps vary in different grid modules along the MEC dorsal-ventral axis. Additionally, our model also predicts theta-modulated head direction cells with specific properties, which we test with empirical data from theta modulated head direction cells recorded in the anteroventral thalamic nucleus [18].

In summary, our model proposes the spontaneous emergence of theta sweeps in hardwired neural networks within the brain’s spatial navigation system. This model enhances our understanding of the generation and nature of theta sequences, potentially shedding light on the neural dynamics underlying various navigational behaviors.

## Results

### The computational model

The computational framework, following that outlined in [13], is composed of continuous attractor networks modeling theta-modulated head direction (HD) cells in the parasubiculum as a ring attractor (HD attractor) and grid cells in the MEC as a two-dimensional attractor on a neuronal sheet (GC attractor) (see Fig. 1a-c, Fig. S1 and Methods for details). When the animal runs in the environment, head direction-dependent sensory input activates an activity bump in the HD attractor. Activated HD cells then activate downstream conjunctive grid x direction cells in MEC layer 3, which have the same preferred firing direction as those activated HD cells. The conj-grid cells further carry a position signal as a phase input onto the GC attractor manifold, with the input strength modulated by the animal’s running speed [19]. Importantly, this phase input is slightly shifted along the internal direction encoded both in the HD cells and conj-grid cells [13], driving the activity bump in the GC attractor (representing the internal location of the animal) to move in that direction.

Neurons in the model exhibit firing rate adaptation. At the single neuron level, this leads to a reduction in a neuron’s firing rate in response to an input of constant intensity (Fig. 1d&e). At the network level, it induces intrinsic mobility of the activity bump by destabilizing it [20]. This intrinsic mobility interacts with the external drive of sensory input, resulting in sweeps of the activity bump [17] (see below).

Furthermore, both the HD cells and the grid cells in the model receive theta rhythmic input from the medial septum [21] with input strength increasing with the animal’s running speed (see Methods for more details). This theta input governs the sweep frequency of the activity bumps induced by firing rate adaptation. In summary, the combination of internal firing rate adaptation and external theta modulation results in internal direction sweeps in the head direction network, which further drives the internal location sweeps in the GC attractor network via an intermediate layer of conj-grid cells (see Methods for more details).

### Bidirectional and forward sweeps in theta-modulated HD cells

We define the location of the activity bump on the HD attractor manifold as the internal direction of the animal [13] (Fig. 2a). We first demonstrate that, with firing rate adaptation, theta-modulated head direction cells exhibit sweeps of internal direction. Depending on whether the animal rotates its head, the internal direction exhibits either bidirectional sweeps or forward-directed sweeps. Specifically, when the animal runs along a straight trajectory, with a fixed head direction, the internal direction exhibits bidirectional sweeps from side to side of the head axis (Fig. 2b). This aligns with the discrete, alternating activity observed from side to side of the head axis in head direction cells during fast and straight running [13].

**Figure 2:**
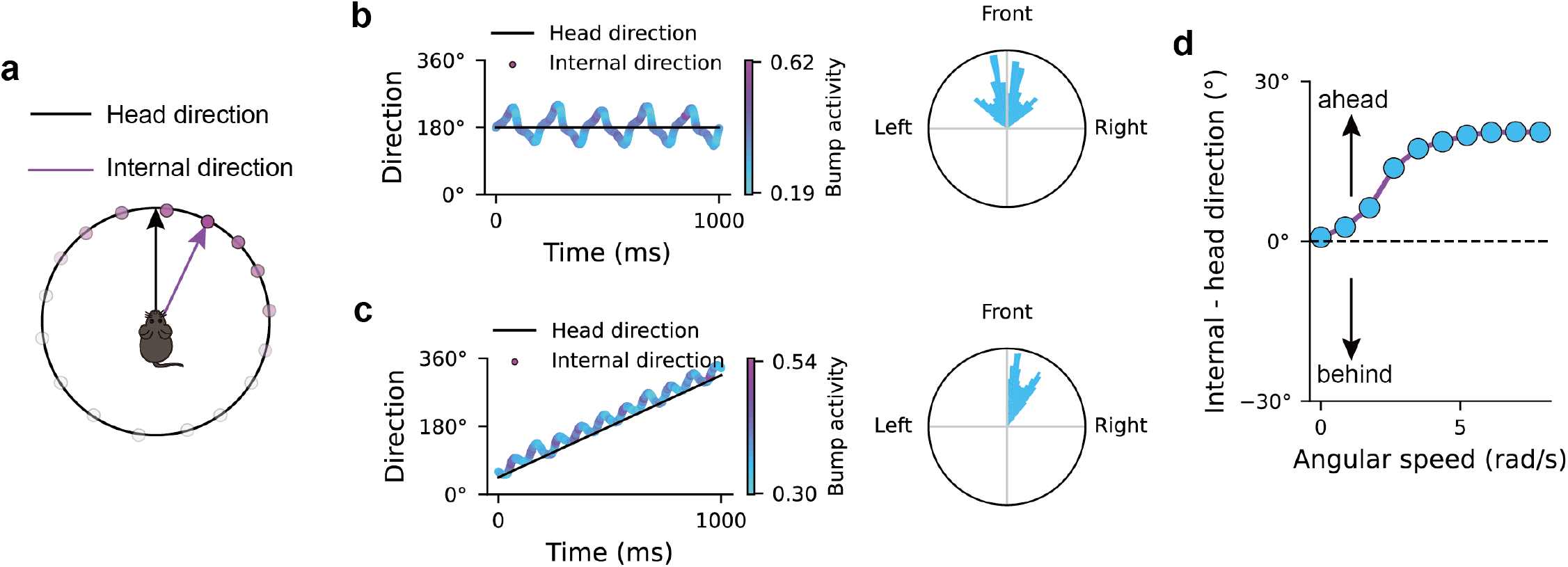
Forward-direct theta sweeps and bi-directional theta sweeps in the HD attractor. **a**. A schematic of head direction and internal direction represented by the activity bump in the HD attractor. **b**. Left: bidirectional theta sweeps when the animal runs along a straight trajectory with fixed head direction (the dark line). Blue to purple dots represent the internal direction (the bump center) across 1-second simulation. More purple dots represent higher peak firing rate of the activity bump. Right: polar histogram of the offset angle of the internal direction relative to the head axis over time. **c**. Left: Forward-direct theta sweeps when the animal turns right (clockwise) with a constant angular speed. Right: polar histogram of the offset angle of the internal direction relative to the head axis over time. **d**.The averaged offset of the internal direction relative to the head axis as a function of head rotation speed. Positive values represent the internal direction leading ahead of the head direction.

Furthermore, when the animal’s head rotates, the internal direction exhibits forward-directed sweeps ahead of the head direction (Fig. 2c). This ahead representation explains previous findings related to anticipatory firing in head direction cells [22, 23]. Importantly, the forward-directed theta sweeps predict the existence of theta phase precession in theta-modulated HD cells (see Results. below). Moreover, we reveal a continuous spectrum from bidirectional sweeps to forward-directed sweeps as the angular speed increases. Specifically, as the animal rotates its head faster, internal direction sweeps become more forward directed in a linear manner (Fig. 2d). While alternating activity in head direction cells is most significantly observed during fast straight runs in open-field environments [13], forward-directed sweeps during turning periods are an important contribution to the overall pattern of results (Fig. S2 and Video.1).

### Internal direction sweeps in HD cells drive alternating location sweeps in grid cells

Here we demonstrate that when the animal runs in a straight line, the bidirectional sweeps of internal direction, combined with firing rate adaptation in the grid cell network, drive left-right alternating sweeps of internal location in grid cells (Fig. 3a&b). Moreover, since the effect of firing rate adaptation in the HD network varies with turning and hence affect the dynamics in downstream grid cells, we also demonstrate theta sweeps in a segment of the real trajectory of a rat, including straight runs, turns and immobile periods (see Fig. 3c and Video.1). For straight runs, the sweep angle of internal location is 24.6 ± 1.3 degrees to either side of the animal’s head axis, while the sweep angle of internal direction is 17.0 ± 0.7 degrees to either side of the animal’s head axis (Fig. 3d), which is comparable to empirical data [13] (see Table. 1 for parameter settings). Importantly, the internal direction sweep in the upstream head direction cells drives the internal location sweep, directing it to the same side of the head axis across theta cycles (see Fig. 3d&e and Vollan et al. [13]). Moreover, the sweep angle of the internal direction contributes to the sweep angle of the internal location (Fig. 3f).

**Figure 3:**
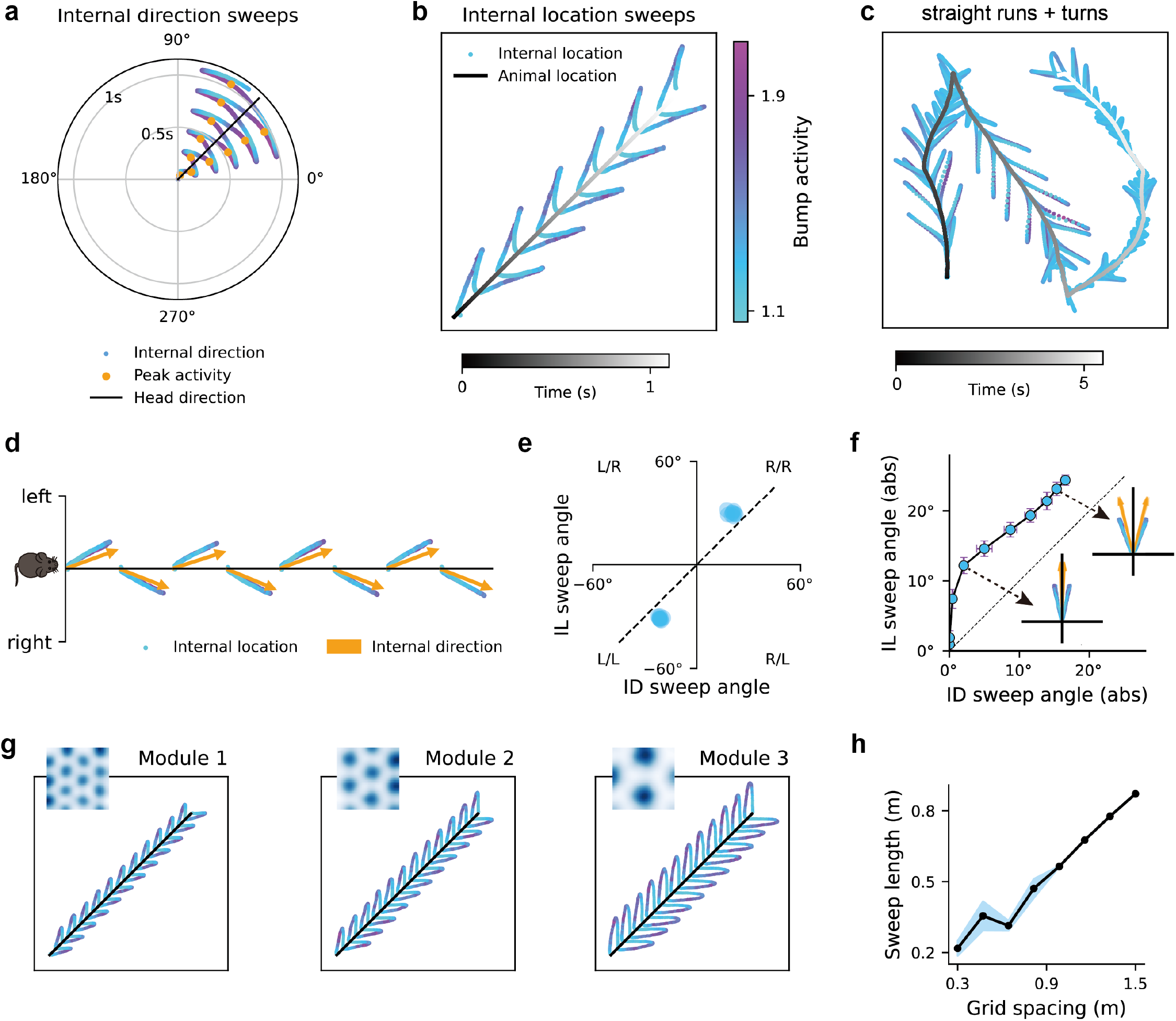
Internal direction sweeps drive left-right-alternating sweeps of internal location in grid cells. **a**. Internal direction sweeps in the model when the animal runs along a straight line with a fixed head direction (45^°^). More purple dots represent higher firing rate of HD cell population. **b**. Internal location sweeps from the pure grid cells with the grey line represents the animal’s trajectory over time and the dots represent the internal location. More purple dots indicates higher firing rate of grid cell population. **c**. Internal location sweeps on a trajectory including straight runs, turns and immobile periods (see also Video.1). **d**. The internal location sweeps in grid cells align with the internal direction sweeps in upstream head direction cells. **e**. A scatter plot of the internal direction sweep angle and the internal location sweep angle across all theta cycles on a simulated straight run. The gray dashed line represents identical value of the two sweep angles. **f**. The internal location sweep angle increases with the internal direction sweep angle, with the insets showing two example sweeps with different angles. The dashed line marks the identical sweep angles of the internal direction and the internal location. **g**. Internal location sweeps in grid modules with different grid spacing. **h**. Location sweep length is proportional to grid spacing. Dots represent the mean of 10 simulations at each grid spacing, with the shaded area representing the standard deviation.

Different grid modules form independent continuous attractor networks [24] and sweep length was found to increase proportionally with the grid spacing in each module [13]. Our model implemented this finding by adjusting the ratio of mapping between the physical space and the phase space defined by the toroidal unit (see Methods). An increased mapping ratio results in larger grid spacing (Fig. 3g). While different attractor networks receive the same speed input in physical space (the animal’s speed), networks with larger grid spacing have proportionally lower speed input in phase space after transformation. Importantly, the speed variation in phase space has only a minor effect on the sweep length on the toroidal manifold (Fig. S3), indicating that the sweep lengths are equal across grid modules when mapped onto the module’s toroidal unit tile [13]. The invariance of sweep length across modules, in turn, when mapped back to physical space, increases proportionally with the mapping ratio, thus correlating linearly with the grid spacing (see Fig. 3g&h and Video.2).

Instead of teleporting instantaneously back to the animal’s location at the end of each theta cycle, the activity bump sweeps continuously back to the animal’s location, because of the continuous nature of the CG attractor. During outward sweeps, grid cells exhibit higher firing rates than during inward sweeps (see Fig. 3b&g and Fig. S4), because the firing of cells on the outward sweep path causes firing rate adaptation, reducing their firing rate when the activity bump sweeps inward. We found the same pattern in experimental data from grid cells (Video.3) notwithstanding the presentation in [13] which emphasizes the forward sweeps.

### Speed modulation of theta sweep features

In our model, running speed increases both the strength of medial-septal theta modulation (consistent with experimental data [25]) and the strength of shifted phase input from the conj-grid cells to the GC attractor network [19]. These two factors jointly affect both the sweep features in the HD attractor network and the downstream GC attractor network (Fig. 4a). We checked four sweep features, including the alternation score (Methods), the mean sweep angle, the variance of the sweep angle, and the internal location sweep length (Fig. 4b). We first verified that the alternation score of theta sweeps in the GC attractor network increases with running speed (Fig. 4c), while the variance of sweep angle decreases with running speed (Fig. 4d). These two results agree with the empirical observation of increased sweep bimodality with running speed [13].

**Figure 4:**
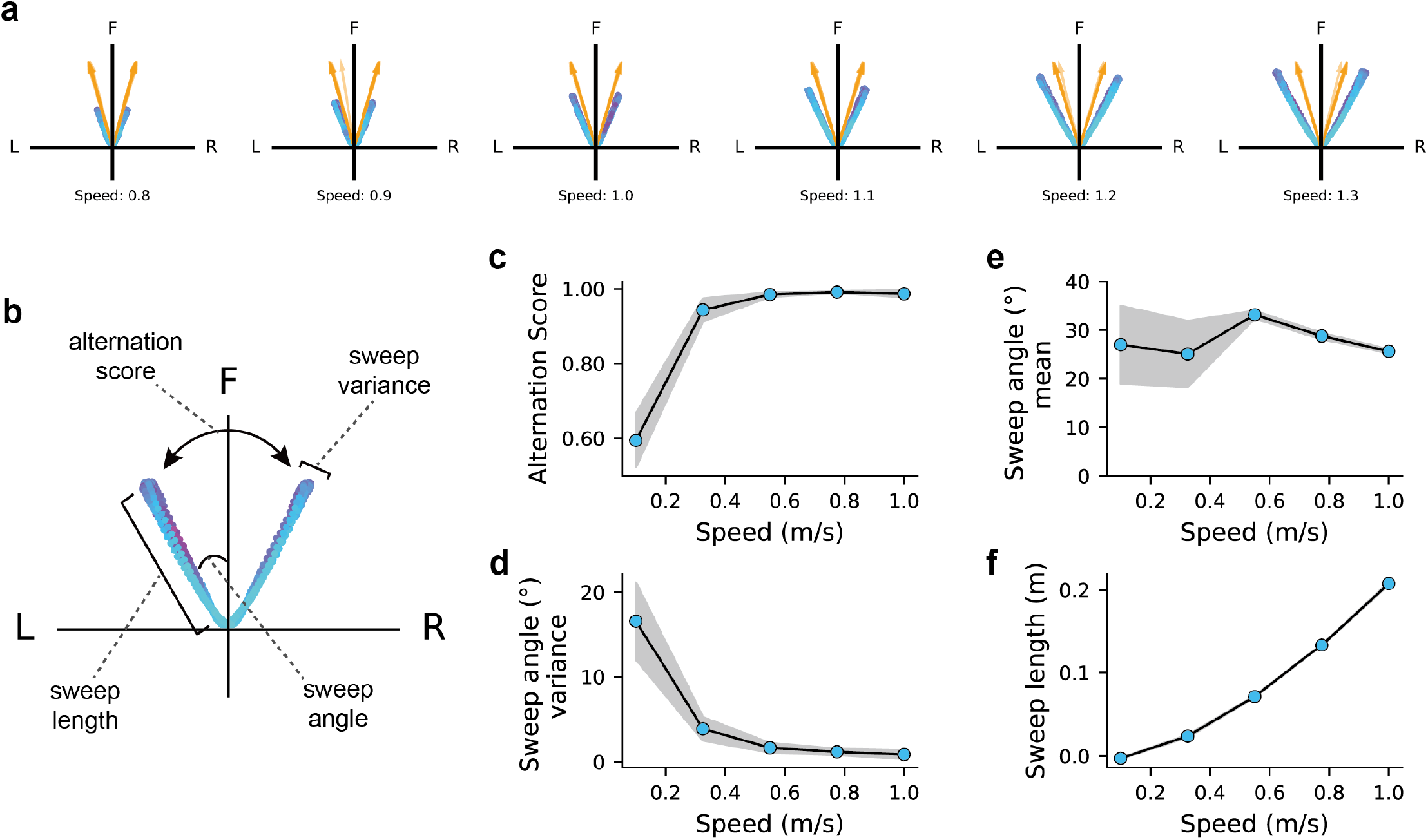
Speed modulation of sweep features. **a**.Left-right alternating theta sweeps of internal direction in the HD attractor (orange arrows) and internal location in the GC attractor (blue-purple dots) under different running speeds. **b**. Schematic of extracting the mean sweep angle, the variance of sweep angle, the sweep length and alternation score. **c**. The relationship between the alternation score of location sweeps in grid cells and running speed of the animal. Shaded area represents the standard deviation of the results from 10 simulations of the model under each running speed. **d**. The relationship between the variance of sweep angle and running speed. **e**.The relationship between the mean sweep angle and running speed. **f**.The relationship between the sweep length and running speed.

We made additional testable predictions related to other sweep features. First, the mean sweep angle remains roughly the same as running speed increases (Fig. 4e), indicating that the sweep angle is minimally affected by running speed. Second, the sweep length increases almost linearly as running speed increases (Fig. 4f). This is also due to stronger input from conj-grid cells at higher speeds [19], which drives the activity bump to move further. If hippocampal place cell sweeps are inherited from entorhinal grid cells [13], this could also explain the linear relationship of theta sweeps and speed found in the hippocampus [26].

### Firing rate adaptation modulates theta sweeps along the MEC dorsal-ventral axis

In the extreme case, without any firing rate adaptation, the internal direction tracks the animal’s head direction, while the internal location tracks the animal’s position (Fig. S5). In the more general case, as adaptation strength increases, our model generates several predictions that can be tested in future experiments. First, increasing the adaptation strength in head direction cells increases the theta sweep angle of the internal direction (Fig. 5a). Second, increasing the adaptation strength in grid cells increases the theta sweep angle of the internal location (Fig. 5b). Furthermore, while sweeps in grid cells are driven by sweeps in HD cells, the location sweep angle is larger than the direction sweep angle, due to the additional firing rate adaptation in grid cells (see Fig. 5b, also Fig. 3f and Fig. 4e). Increasing the adaptation strength in grid cells increases the angular difference between the location sweep and the direction sweep (Fig. 5b).

**Figure 5:**
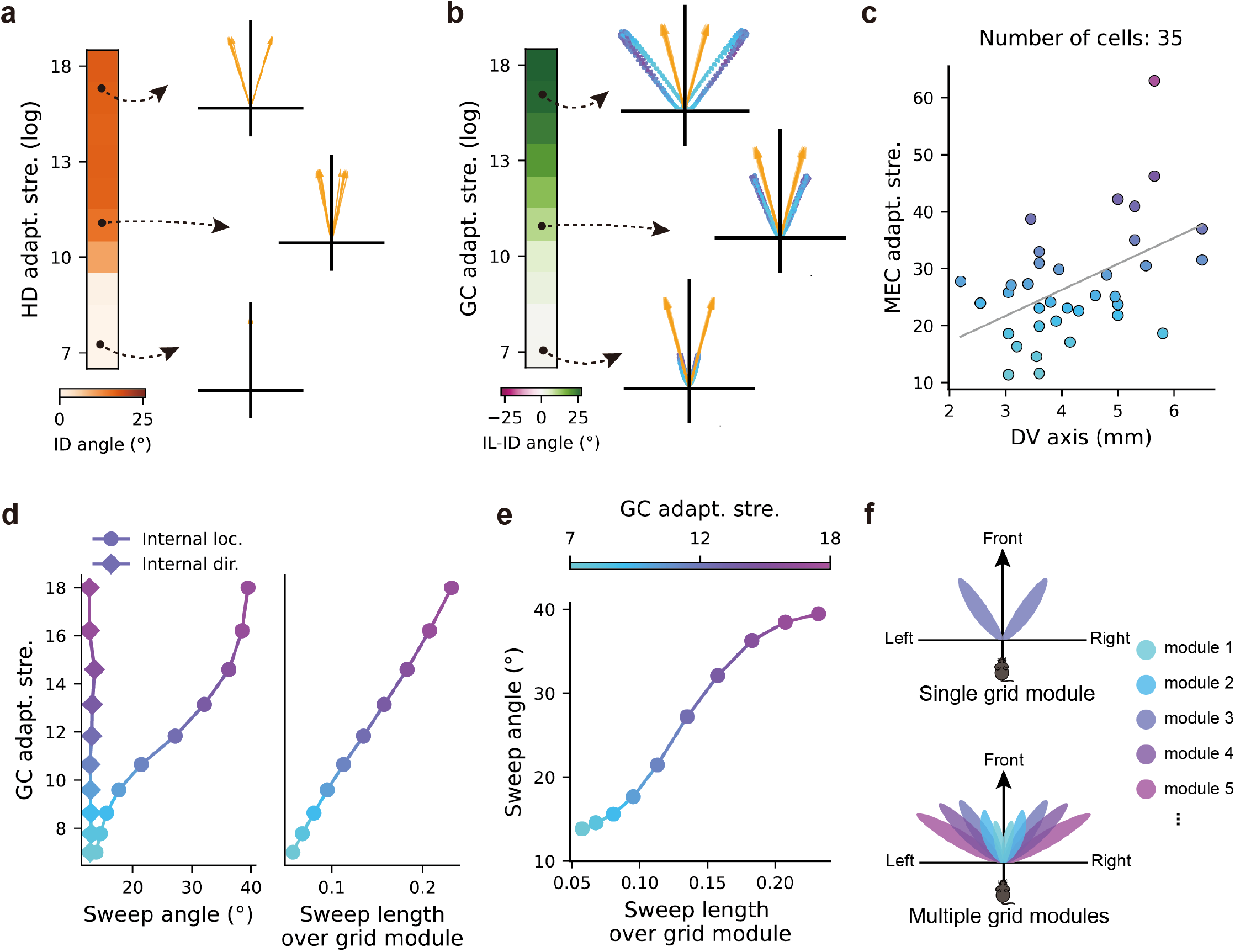
Adaptation strength modulates theta sweeps along the MEC dorsal-ventral axis. **a**. The sweep angle of internal direction increases with the strength of firing rate adaptation in head direction cells, with three examples shown on the right. **b**. The angular difference between the location sweep and the direction sweep increases with the strength of firing rate adaptation in grid cells (while the adaptation strength in head direction cells is fixed), with three examples shown on the right. Green/pink color represents larger/smaller sweep angle in internal location than that in internal direction. Note that the angular difference is always positive, meaning that the location sweep angle is larger than the direction sweep angle. **c**. The adaptation strength as a function of locations along the dorsal-ventral axis for 35 cells in the MEC with in-vitro whole-cell patch clamp recording. The grey line show a linear fit (Pearson correlation with *r* = 0.48, *p* = 0.003). The data was obtained from Yoshida et al. [27] with permission. **d**. Left: the sweep angles as a function of increased adaptation strength in the GC attractor network (with the adaptation strength in the HD attractor network fixed). Right: same as the left but shows the sweep length of internal location (normalized by the effect of grid spacing) as a function of increased adaptation strength along the dorsal-ventral axis. **e**. The sweep angles correlates positively with the sweep lengths (normalized by the effect of grid spacing) while increasing the adaptation strength in grid cells along the dorsal-ventral axis. **f**. Schematic of sampling of surrounding space with a single grid module versus with multiple grid modules. More purple colors represents sweeps in grid modules with larger adaptation strength and larger grid spacing.

While evidence for changes of adaptation strength across head direction cells is lacking, the strength of firing rate adaptation increases along the MEC dorsoventral axis [27] (Fig. 5b). This leads to a third prediction, that the location sweep angle is larger in grid modules located in the more ventral areas of the MEC (Fig. 5c&d). Moreover, stronger adaptation also leads to an increase of the sweep length of internal location, even when expressed as a proportion of grid scale (Fig. 5d). This suggests a positive correlation between sweep angle and sweep length across multiple grid modules along the dorsal-ventral axis because of the increase in adaptation strength (Fig. 5e); that is, shorter sweeps in dorsal grid modules with smaller grid spacing are more forward-directed, while longer sweeps in ventral grid modules with larger grid spacing alternate more left-to-right. When the animal runs in an open field, theta sweeps from multiple grid modules can therefore cover a much larger surrounding space than single-module theta sweeps, potentially contributing to efficient sampling of the surrounding space without the need for physically visiting it (Fig. 5f).

### Theta phase precession correlates with theta skipping in head direction cells

While classic HD cells have been reported in many brain regions such as anterior thalamus, lateral mammillary bodies and retrosplenial cortex, theta modulated HD cells have also been found recently in the anteroventral thalamic nucleus [28, 18], parasubiculum and entorhinal cortex [29, 13]. By varying the theta input and firing rate adaptation, our HD attractor module generates three types of HD cells observed in previous studies (Fig. 6a-c and Fig. 7a&b). Specifically, 1) without medial-septal theta input, cells in the HD attractor exhibit the features of classic HD cells (Fig. 6a). The activity bump leads ahead of the animal’s head direction when rotating, giving rise to anticipatory firing in HD cells [22, 23]; 2) with theta input but without firing rate adaptation, cells in the HD attractor exhibit the features of theta modulated HD cells where HD cells fire in every theta cycle (Fig. 6b); 3) with both theta input and firing rate adaptation, cells in the HD attractor exhibit the features of theta skipping HD cells where HD cells fire in alternating theta cycles (Fig. 6c) [30, 29].

**Figure 6:**
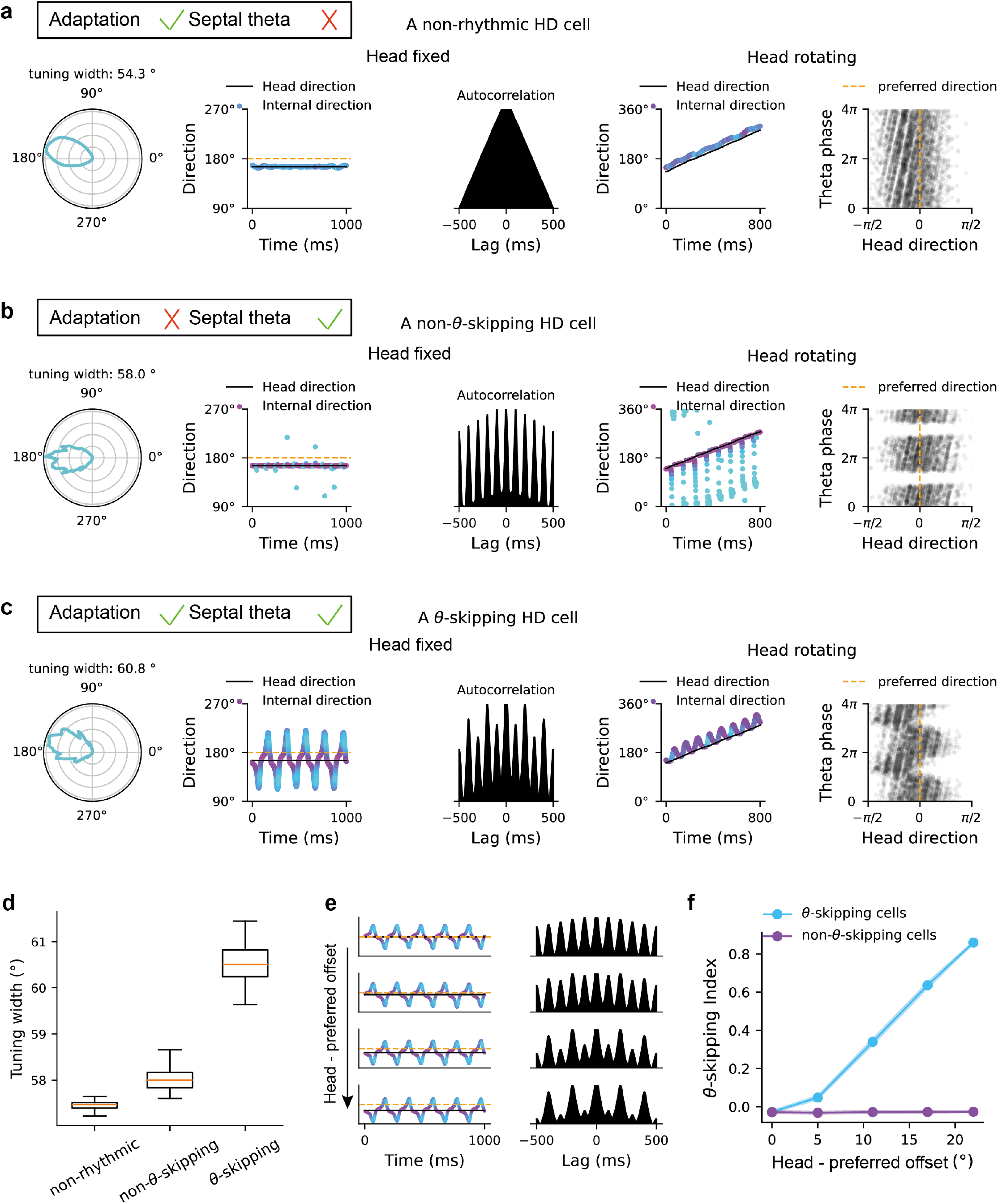
Theta skipping and theta phase precession in the head-direction ring attractor. **a**. A classic HD cell without theta modulation, simulated with a HD attractor with firing rate adaptation but without septal theta input. From left to right: the tuning field of a cell; the internal direction over 1 second time window when the animal’s head is fixed, with orange line representing the preferred direction of a probed cell; the auto-correlation of the firing of the probed cell; the internal direction over 1 second time window when the animal’s head is rotating with a constant angular speed; the spike phase of the probed cell as a function of head direction. **b**. A HD cell with theta modulation but without theta skipping, simulated with septal theta input but without firing rate adaptation. **c**. A HD cell with theta skipping, simulated with both septal theta input and firing rate adaptation. **d**. The tuning width of the three types of HD cells. **e**. Increased theta skipping effect as the fixed head direction (dark lines) is more away from the preferred direction of a probed cell (orange lines). **f**. Theta skipping index increases as a function of the offset between the head direction and the preferred direction of a cell for theta skipping HD cells (blue), but not for normal theta-tuned HD cells without skipping (purple).

**Figure 7:**
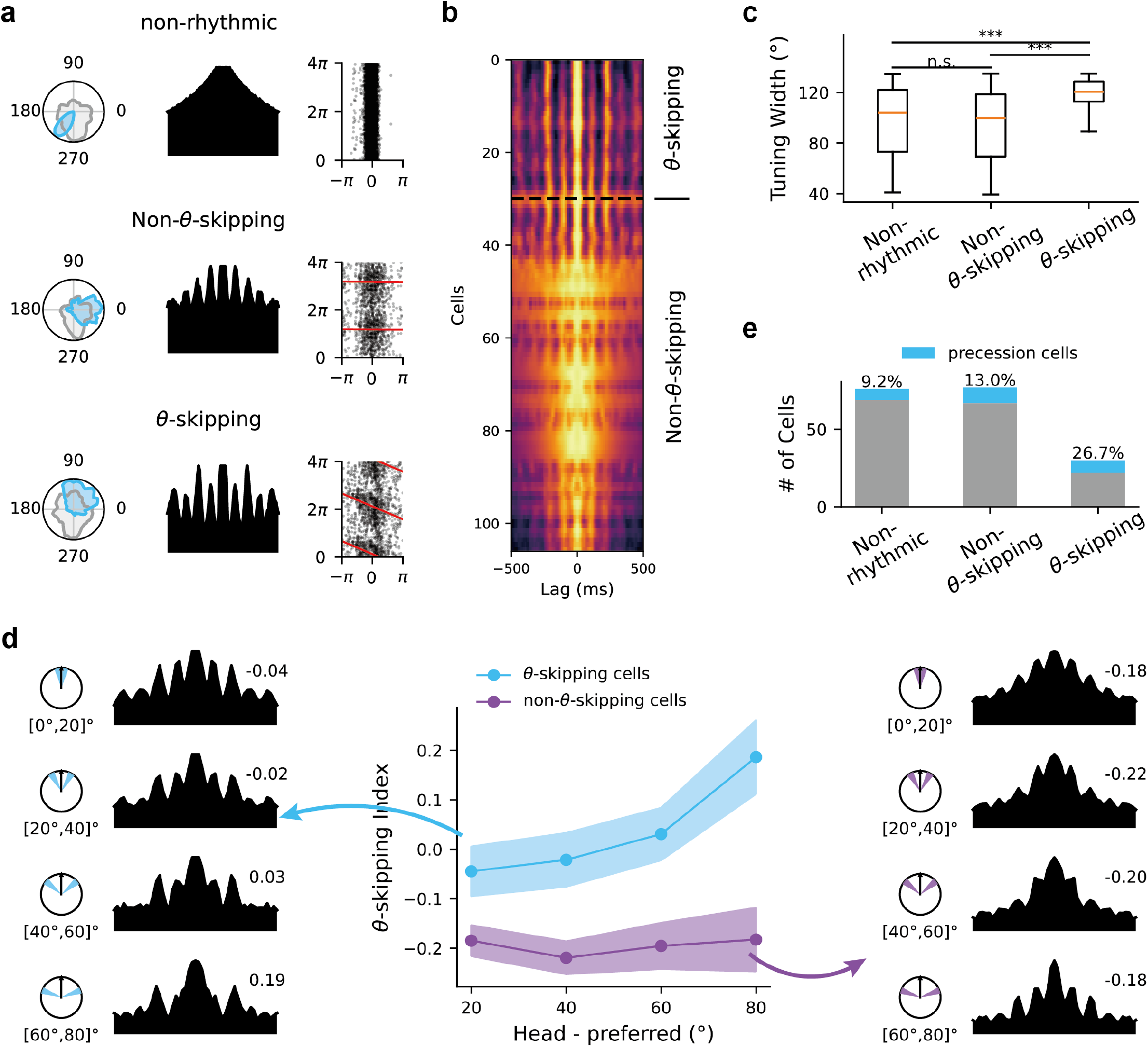
Theta skipping and theta phase precession in the HD cells in the anteroventral thalamic nucleus. **a**. Examples of a classic HD cell (non-rhythmic), a normal theta-tuned HD cell without skipping (non-theta-skipping) and a theta skipping HD cell. From left to right: the tuning width, the auto-correlation, the spike phase versus head direction with preferred direction realigned to 0. Red lines represent circular-linear fit. **b**. Heat map of auto-correlations of all theta modulated HD cells, ordered by theta skipping index. Warmer colors represent higher correlation values. Top rows are for theta skipping cells and bottom rows are for normal theta-tuned HD cell without skipping. **c**. The tuning width of the three types of HD cells. **d**. Theta skipping index as a function of the offset between head direction and the preferred direction for both non-theta-skipping HD cells (purple) and theta skipping HD cells (blue). The auto-correlation plots were taken by averaging the auto-correlation of each cell in the offset range showed on the disk, with preferred directions all aligned to the north. **e**. The percentage of cells showing significant phase precession (with the circular-linear correlation *p* < 0.05) in three types of HD cells.

From the populational view, theta skipping of individual neurons can originate from the sweeps of activity bumps, which leads to two interesting predictions. First, bump sweeps increase the tuning width of theta skipping HD cells compared to classic HD cells and normal theta modulated HD cells where there is no bump sweep (Fig. 6d). Second, the theta skipping effect becomes more significant when the animal runs along a direction with a larger offset to the preferred direction of the cell, as the activity bump is more likely to sweep over the preferred direction in alternating theta cycles (Fig. 6e&f).

Finally, our HD attractor with firing rate adaptation and medial-septal theta input predicts the existence of theta phase precession in theta skipping HD cells when the animal turns through the preferred direction of the cell (Fig. 6c). This corresponds to network models of theta phase precession in place cells when the animal runs in linear track environments [14, 15, 17], where the activity bump sweeps over the preferred location of a place cell at progressively earlier phases of theta cycles when the animal traverses the firing field.

We verified the three predictions related to directional tuning width, theta skipping and theta phase precession with empirical data containing a larger number of head direction cells recorded from the anteroventral thalamic nucleus (Fig. 7a&b; number of classic HD cells: 76; normal theta modulated HD cells: 77; theta skipping HD cells: 30). First, directional tuning width is significantly larger in theta skipping HD cells than normal theta modulated HD cells (Mann-Whitney U Test [MWUT] with *Z* = 4.6, *p* = 5.1 × 10^−6^) and classic HD cells ([MWUT] with *Z* = 3.8, *p* = 1.5 × 10^−4^). However, the tuning width in normal theta modulated HD cells and classic HD cells are similar ([MWUT] with *Z* = 1.0, *p* = 0.306) which agrees with the model where the activity bump does not sweep in these two cases. Second, for theta skipping HD cells, the skipping effect increases during the periods when the animal’s head direction is more away from the preferred direction of the cell, while for normal theta modulated HD cells, the skipping effect remains weak and independent of the animal’s head direction (Fig. 7d). Third, some HD cells recorded in the anteroventral thalamic nucleus show significant theta phase precession during turning. Moreover, the percentage of phase-precession HD cells is higher in theta skipping HD cells than that in normal theta modulated cells and classic HD cells (Fig. 7e; classic HD cells: 8/76, 9.2%; normal theta modulated HD cells: 10/77, 13.0%; theta skipping HD cells: 7/30, 26.7%).

### Theta sweeps disappear with septal inactivation

Reduction in theta oscillation by pharmacological inactivation of the medial septum correlates with impairment in spatial memory tasks [31]. At the neurophysiological level, septal inactivation eliminates theta skipping in HD cells in the parasubiculum and MEC [29], and impairs phase precession in the entorhinal cortex and hippocampus [32]. Inspired by these phenomena, we investigated how septal inactivation affects network dynamics in our model.

First, we showed that in the head direction network (Fig. 8a), reducing the theta rhythm magnitude decreases the occurrence of direction sweeps and therefore theta skipping in HD cells (Fig. 8b&c), which aligns with previous empirical results [29]. Moreover, since theta skipping and phase precession are two aspects of the same network (depending on whether or not the head is rotating), we predicted that reducing the theta rhythm magnitude also decreases the effect of theta phase precession in HD cells (Fig. 8b&d).

**Figure 8:**
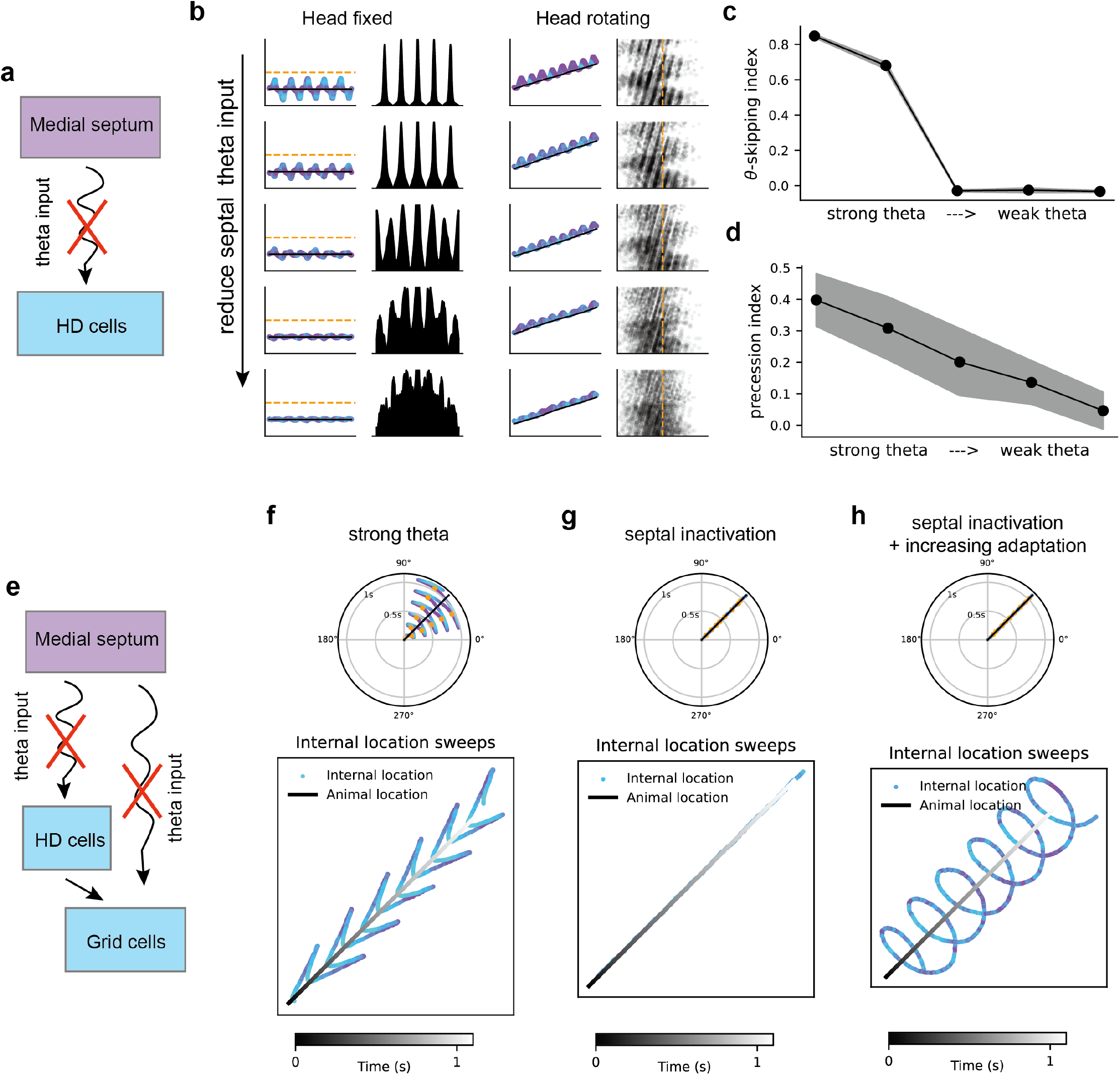
Theta sweeps disappear with septal inactivation. **a**. Schematic of inactivating medial-septal theta input to the HD cells. **b**. Reduced effect of theta skipping with head fixing (left two columns) and theta phase precession with head rotating (right two columns) as weakening the theta input strength from the medial septum. **c**. Theta skipping index decreases as weakening the theta input strength. **d**. Circular linear correlation coefficient decreases as weakening the theta input strength. **e**. Schematic of inactivating medial-septal theta input to both the HD cells and the grid cells. **f**. left-right sweeps in grid cells with medial-septal theta input. **g**. The activity bump tracks the animal’s position without medial-septal theta input. h. Circular sweeps in grid cells without medial-septal theta input under increased adaptation strength.

Second, we showed that in the downstream grid cell network (Fig. 8e), inactivating the septal theta rhythm leads to two dynamics different from the left-right sweeps under normal settings of firing rate adaptation and septal theta input (Fig. 8f). Specifically, 1) with relatively weak adaptation in grid cells, both the internal direction and location track the animal’s head direction and position (Fig. 8g); 2) with relatively strong adaptation in grid cells, while the internal direction still tracks the animal’s head direction, the internal location exhibits circular sweeps around the animal’s position for 360^°^ within individual theta cycles (Fig. 8h). While such circular sweeps have not been reported in empirical data, they might exist as intrinsic network dynamics under weak theta modulation from the medial septum when running speed is low.

These predictions could be tested in future inactivation experiments, bearing in mind that rotational theta may have a cholinergic component [33, 34] compared to translational theta which is controlled by GABAergic and glutamatergic medial septal cells [35].

## Discussion

We demonstrated that a continuous attractor model with firing rate adaptation and medial-septal theta modulation is sufficient to explain recent experimental findings of left-right-alternating sweeps in grid cells and related directional “flickering” in theta-modulated head-direction cells Vollan et al. [13]. In this model, the upstream head direction network generates internal direction sweeps at the theta rhythm, which activates the conjunctive grid x direction cells with preferred direction aligning with the internal direction. These conj-grid cells then provide a shifted input along the same direction, driving the left-right alternating sweeps in grid cells in a coordinated manner. This model accounts for many of the experimental findings, including the alignment of internal direction sweeps in parasubicular HD cells and internal location sweeps in grid cells, with the former driving the later; the linear relationship between location sweep length and grid spacing across different grid modules; and the increase of the sweep alternation score with running speed.

The proposed model improves in two aspects over previous network-based models of theta sweeps and theta phase precession when the animal runs along a linear track [14, 15, 16, 17]. First, while phase precession has been observed in both grid cells and place cells in open fields [36, 37], models for populational theta sweeps extending beyond linear track scenarios have not been well addressed so far. However, recent progress has been made, including our modeling of left-right theta sweeps in place cell networks when the animal runs toward a decision point in a T-maze [17, 12], as well as a similar model with firing rate adaptation and theta input that accounts for left-right alternating sweeps in place cells in an open field [38]. Second, we systematically modeled theta sweeps in a system of HD cells and grid cells, with sweep direction in HD cells determining sweep direction in downstream grid cells and place cells [13]. Instead of generating theta sweeps independently in grid cell networks across multiple grid modules and place cell networks, the coordination of sweep directions across multiple networks by head direction cells may help synchronize the temporal organization of spiking activity across multiple brain regions. This synchronization could potentially support the stabilization of a single coherent map for spatial navigation and episodic memory function [39, 40, 41].

The model only requires attractor dynamics, firing rate adaptation and theta rhythmicity to generate coherent theta sweeps. Sensory input or actual running is not strictly necessary. In our simulations the theta input increases with running speed, but a strong theta input on its own (without a moving bump of sensory input) will still generate theta sweeps, as seen during R.E.M. sleep [13]. Equally, the decoding of theta sweeps occurs on the toroidal manifold of the grid attractor, and is then mapped onto physical space with a consistency constraint to resolve periodic ambiguity (see Methods), which allows decoding of theta sweeps to unvisited locations [13].

Our modeling and empirical results favor continuous sweeps instead of discrete flickers from side to side of the animal’s head axis in the HD network and forward only grid cell sweeps as presented by Vollan et al. [13]. These continuous sweeps align with the continuous-attractor nature of the HD system [42, 43] and our finding of theta phase precession during turning, and may also affect the continuity of theta sweeps in downstream grid cells. Using data from Gardner et al. [24], we decoded theta sweeps from grid cell population activity and found that the decoded position likely sweeps continuously backward to the animal’s location within individual theta cycles (see Fig. S6 and Video.3). While our model favors continuous sweeps in grid cells, we did observe higher network activity during outward sweeps compared to inward sweeps (see Fig. 2b&c, Fig. 3a and Fig. S4, also see [14, 15]), which may affect the continuity of decoded position within each cycle. Thus, we consider the continuous sweeps reported here, compared to the left/right “flickering” of directional sweep and forward-only locational sweeps reported by Vollan et al. [13], to reflect a quantitative difference in thresholding for presentation rather than a real qualitative difference.

Our model makes numerous novel predictions related to the features of theta sweeps in both head direction cells and grid cells. In theta-modulated head direction cells, we predicted: 1) increased directional tuning width in theta skipping cells; 2) increased theta skipping effect when the offset between the head direction and the preferred direction increases; 3) the existence of theta phase precession in theta skipping cells; all of which we verified with empirical data from theta-modulated head direction cells in anteroventral thalamus [18].

In grid cells, we predicted: 1) as the animal runs faster, the sweep length increases while the variance of the sweep angle decreases; 2) within a single grid module, the sweep angle in grid cells is always larger than that in upstream head direction cells; 3) across multiple grid modules along the MEC dorsal-ventral axis, there is a systematic increase in both the length and angle of theta sweeps. Compared to single-module sweeps, the multi-module sweeps provide an efficient way of covering a surrounding space without physically visiting them, potentially contributing to an efficient formation of new maps for spatial navigation (Fig. 5e&f). Moreover, we predicted reduced effects of theta skipping, phase precession and theta sweeps when the medial-septal theta rhythm magnitude decreases, which could be tested with septal inactivation in future experiments.

Our model suggests that two temporal firing features in theta-modulated HD cells, i.e., theta skipping and theta phase precession, are two aspects of the same underlying network dynamics of theta sweeps. This is supported by two empirical observations with data recorded in the anteroventral thalamic nucleus [18]: 1) cells exhibiting theta skipping have a higher probability of showing theta phase precession (Fig. 7e); 2) theta skipping HD cells, but not normal theta modulated HD cells, show an increase in the skipping index as the animal’s head direction deviates further from the cells’ preferred directions (Fig. 7d). Since the classic head direction signal is generated subcortically, arriving to neocortex via the mammillary bodies and anterodorsal thalamus at the dorsal presubiculum [44], we hypothesize that theta-modulated HD cells in the parasubiculum inherit their firing properties from the anteroventral thalamic nucleus via the presubiculum [45]. Although some studies show that HD cells in the dorsal presubiculum are not theta-modulated [46, 47], others reported theta-modulated HD cells in more ventral presubiculum [48, 49]. A clarification of the circuit for theta-modulated HD cells in future experiments would be welcome.

In summary, we propose a simple yet comprehensive network model where left-right-alternating theta sweeps can be generated intrinsically during open field navigation. This theoretical framework aligns with numerous experimental findings and provides testable predictions for future research. These results enhance our understanding of the neural dynamics underlying spatial navigation and memory-related cognitive functions.

## Acknowledgments

We thank Richard Gardner and Edvard Moser, and Eleonora Lomi and Kate Jeffery and their research groups for making their grid cell and head direction cell data publicly available online. We thank Motoharu Yoshida and Michael Hasselmo for sharing their whole cell patch clamp data. The work was supported by a Wellcome Principal Research Fellowship (222457/Z/21/Z, N.B.), and a Science and Technology Innovation 2030-Brain Science and Brain-inspired Intelligence Project (2021ZD0200204, S.W.).

## Author contributions

All authors conceptualized the project. ZJ and TC built the model and performed the simulations with supervision from NB and SW. ZJ analyzed the experimental data with input from NB. All authors discussed and interpreted the results. ZJ and NB wrote the paper.

## Code and data availability

Code for reproducing all the results (including modeling and experimental data analysis) will be available before publication. The grid cell dataset is from Gardner et al. [24] and available at: https://figshare.com/articles/dataset/Toroidal_topology_of_population_activity_in_grid_cells/16764508. The head direction cell dataset is from Lomi et al. [18] and available at: https://figshare.com/articles/dataset/_strong_Data_code_for_Lomi_et_al_2023_strong_/22802861.

## Methods

### Details of the computational model

The computational framework is composed of continuous attractor networks network modeling theta-modulated head direction cells in the parasubiculum as a ring attractor (HD attractor) and grid cells in the MEC as a two-dimensional attractor on a neuronal sheet. The HD attractor provides directional input, which causes a shifted phase input to the grid cells along the direction of the input signal via an intermediate layer of conjunctive head × grid (conj-grid) cells. Below, we introduce each part of the model in details. For parameter settings, see Table. 1&2.

#### The head direction attractor network

Theta modulated HD cells are modeled as a ring attractor with the following form:

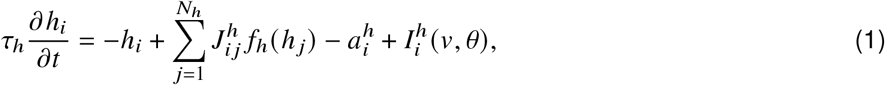

where *h*_*i*_ is the pre-synaptic input to *i*_*th*_ HD cell with *i* = 1, …, *N*_*h*_, and *N*_*h*_ is the number of HD cells. Each HD cell receives recurrent input from other HD cells, along with firing rate adaptation 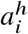 with a form of a negative feedback (see below), and the head-direction dependent sensory input 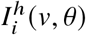. The directional input is modeled with a Gaussian form, with the peak representing the animal’s current head direction, written as,

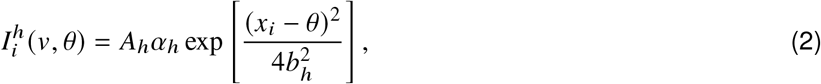

where *α*_*h*_ represents theta modulation from medial septum with a form of sinusoidal wave (see details below). *A*_*h*_ represents the baseline input strength of head-direction cells. *x*_*i*_ is the preferred head-direction of the *i*_*th*_ neurons on the ring attractor, and *b*_*h*_ controls the width of the external Gaussian input. *f*_*h*_ (*h* _*j*_) is the firing rate of the *j*_*th*_ HD cell. The activation function *f*_*h*_ is modeled as a global inhibition, which is written as,

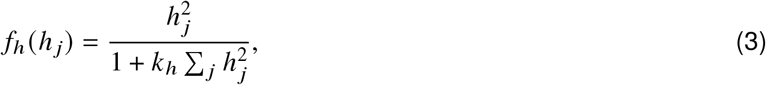

where *k*_*h*_ represents the global inhibition strength among head-direction cells. The global inhibition is important for maintaining a localized activity bump on the ring attractor.

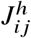 represents the recurrent connections between the HD cell with preferred direction *i* and the HD cell with preferred direction *j*, with the form:

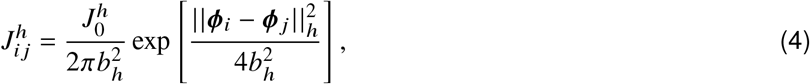

where || · ||_*h*_ denotes the circular distance on the ring and 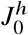 denote the connection strength between cells. *b*_*h*_ controls the tuning width of HD cells.

#### The grid cell attractor network

The pure grid cells are modeled as a two-dimensional continuous attractor network with the following form:

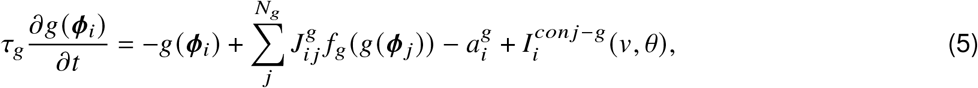

where *g*(***ϕ***_*i*_) is the pre-synaptic input to *i*_*th*_ grid cell with *i* = 1, …, *N*_*g*_, and *N*_*g*_ is the number of grid cells tiling the two-dimensional neuronal sheet. ***ϕ***_*i*_ = (*ϕ*_*x*_, *ϕ*_*y*_) with *ϕ*_*x*_, *ϕ*_*y*_ ∈ (0, 2*π*] representing the preferred phase of the *i*_*th*_ grid cell on the attractor sheet. We set the neuronal sheet with periodic boundaries so that it forms a toroidal manifold. To account for the hexagonal firing fields of grid cells, a mapping function is applied to transform between the location in the physical space and the location in the phase space, which is written as:

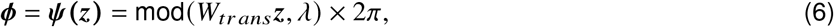

where ***z***(*x, y*) is the coordinate of a location in the physical space, *λ* controls the grid spacing, mod denotes the modulo operation and *ψ* denotes the mapping operator. This mapping takes two steps: first, the transformation matrix *W*_*trans*_ given by:

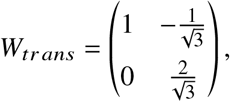

projects ***z*** onto two axes that are sixty degrees apart; second, the modulo operation is applied to convert the transformed coordinate into a phase coordinate. This means when the animal runs into a location away from the current location, the two locations can be wrapped to similar phases in the phase space. This mapping process leads to a grid cell with six-fold symmetry in its firing pattern [50].

Grid cells in different grid modules have different grid spacing, and can operate as independent continuous attractor networks [24]. To model theta sweeps in different grid modules (as shown in Fig. 3g and Video.2), we vary *λ* above to implement them.

*f*_*g*_ (*g*(***ϕ*** _*j*_)) represents the firing rate of the *j*_*th*_ grid cell, which is also modeled as a global inhibition with a form the same as in head direction cells. We denote *k*_*g*_ as the global inhibition strength among grid cells within a module and maintain it with the same value across grid modules. 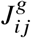 represents the recurrent connections between the grid cell with the preferred phase ***ϕ***_*i*_ and the grid cell with the preferred phase ***ϕ*** _*j*_, with the form:

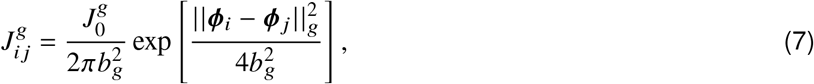

where || · ||_*g*_ denotes the circular distance on the toroidal manifold, and 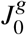 denote the connection strength. *b*_*g*_ controls the grid field width.

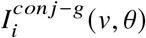 represents the shifted phase input to the *i*_*th*_ grid cell from the upstream conj-grid cells. While we didn’t explicitly model these cells in the current model (due to a high memory consumption since we need to model a 2D attractor at each direction), the shifted phase input has three important features: first, the input strength is modulated by running speed; second, it is also modulated by septal theta oscillation as the conj-grid cells are tuned to theta rhythm [19]. Third, the direction of phase offset is aligned with the internal direction from the upstream HD attractor [13]. Combining all these features and re-writing 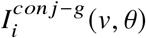 as 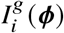 for simplicity, the shifted phase input is expressed as:

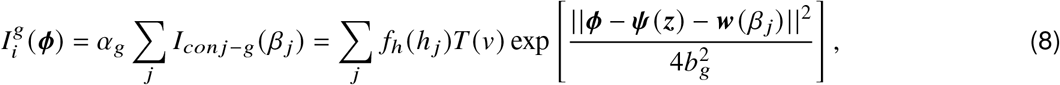

where ***ϕ*** is the phase coordinate, *α*_*g*_ represents theta modulation (see below), *I*_*con j*−*g*_ (*β* _*j*_) is the input from the *j*_*th*_ group of conjunctive grid cells with preferred direction as *β* _*j*_. The resulting phase input equals to a weighted sum of many 2D shifted Gaussian bumps from all the conj-grid cells with the weights as the firing rate of upstream HD cells *f*_*h*_ (*h* _*j*_). ***w***(*β* _*j*_) is a vector: ***w***(*β* _*j*_) = *w*_0_(cos *β* _*j*_, sin *β* _*j*_), where *w*_0_ represents the offset length. This term means if the conj-grid cells coding for the direction of *β* _*j*_ are activated, they will lead to a shifted phase input with the amount of *w*_0_ along the same direction onto downstream grid cells. The strength of the resulting shifted phase input is modulated by running speed with *T* (*v*):

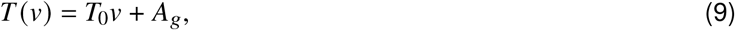

where *T*_0_ and *A*_*g*_ are both constants. This linear modulation reflects that as the animal runs faster, the firing rate of the conj-grid cells increases accordingly [19]. Consequently, a higher speed can lead to a stronger shifted phase input, which drives the activity bump moves faster along the moving direction, and therefore potentially contribute to path integration.

#### Speed modulation of theta input from the medial septum and phase input from the conj-grid cells

In our model, running speed modulates both the oscillation magnitude of medial-septal theta input and the phase input strength from conj-grid cells [19]. Theta modulation onto HD cells is defined as:

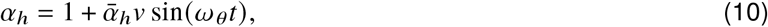

where the oscillation magnitude scales linearly with the running speed *v* [25], with 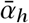 as a scaling factor. Similarly, theta modulation onto grid cells is defined as:

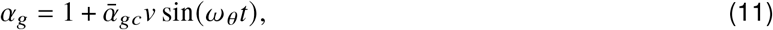

where the oscillation magnitude also scales linearly with the running speed *v*, with 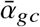 as a scaling factor. While our previous modeling work [17] has demonstrated that firing rate adaptation generates bump sweeps in a continuous attractor model of place cells, the inclusion of medial-septal theta modulation naturally constrains the bump sweep frequency to the theta rhythm, eliminating the need for precise parameter tuning. This addition enhances the model’s biological plausibility and robustness.

#### Firing rate adaptation

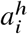 and 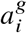 represent firing rate adaptation in the *i*_*th*_ HD cell and the *i*_*th*_ grid cell, both of which have the following form:

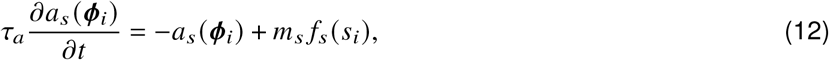

where *s* ∈ {*h, g*}, and *f*_*s*_ (*s*_*i*_) represents the firing rate of the *i*_*th*_ cell. *τ*_*a*_ is the time constant of firing rate adaptation, which is much larger compared to *τ*_*h*_ and *τ*_*g*_, indicating that the process of firing rate adaptation is a slow dynamics compared to that of cell firing. *m*_*s*_ represents the adaptation strength, which can be different in HD cells and grid cells, as well as in different grid modules along the dorsal-ventral axis [27]. At single neuron level, firing rate adaptation reduces the firing frequency following an initial increase in response to an input of constant intensity. At the network level, it induces intrinsic mobility of the activity bump. In fact, any forms of slow negative feedback could exhibit similar effect of increasing the intrinsic mobility, i.e., short-term depression [15], and hence lead to similar network-level dynamics presented in the current study.

### Decoding theta sweeps in our model from the toroidal manifold to the physical space

To decode the internal location in the physical space in our model (see below for the decoding of location in real data), we take the location corresponding to the center of mass of the activity bump in the phase space of the grid cell torus and then transform it back to physical space using the inverse of the transformation matrix. While the mapping from physical location to phase location is one-to-one, the mapping from phase location to physical location is one-to-many. Therefore, to determine which physical location it maps to, we choose the one at the beginning of the simulation that is closest to the animal’s actual location. This approach reduces the one-to-many mapping to a one-to-one mapping at the start of the simulation and allows continuous one-to-one decoding from phase location to physical location until the end of the simulation. Since the phase location on the toroidal manifold does not have a boundary effect, this inverse mapping can result in a decoded position outside the current environment boundary, as reflected in Video.1 and Vollan et al. [13].

### Experimental data

**The grid cell dataset** was collected by Gardner et al. [24] from three experimentally naive male Long Evans rats (Rats 1, 2 and 3) as they foraged for randomly scattered food crumbs in a 1.5 m × 1.5 m square open-field arena. Animals’ 2D position and head direction were obtained from 3D motion capture by attaching a set of five retroreflective markers to implant during recordings. Neural recordings were obtained via Neuropixels silicon probes targeting the MEC–parasubiculum region (one recording session for Rat 1 with single-shank probes in bilateral hemispheres, one recording session for Rat 3 with multiple-shank probes in the left hemisphere, and two recording sessions for Rat 2 with single-shank probes in bilateral hemispheres). Grid cells were identified with methods described in Sargolini et al. [19], which resulted in a total of 1330 grid cells. Specifically, for Rat 1, there were 163 grid cells from two grid modules; for Rat 2, there were 483 grid cells from three modules in session 1 and 544 grid cells from three grid modules in session 2; for Rat 3, there were 140 grid cells from one grid module. We decoded theta sweeps from Rats 1 and 2 since decoding of location was most reliable due to cells were recorded from multiple grid modules (see Fig. S6 and Video.3). This dataset is publicly available at https://figshare.com/articles/dataset/Toroidal_topology_of_population_activity_in_grid_cells/16764508.

**The head direction cell dataset** was collected by Lomi et al. [18] from six adult male Lister Hooded rats as they foraged for randomly scattered food in a 0.9 m × 0.9 m square arena. Animals’ 2D position and head direction were obtained from video tracking of two light-emitting diodes (LEDs) on the headstage. Neural recordings were obtained via chronic recording electrodes targeting the anteroventral thalamic nucleus. Head direction cells were identified using the Rayleigh vector (passing 99^*th*^ percentile shuffle cutoff). To further identify theta modulated HD cells, Lomi et al. [18] computed the index of rhythmicity (IR) and the index of theta phase coupling (IC) for each cell. A HD cell passing 99^*th*^ percentile shuffle cutoff for IC and having an IR ≥ 0.001 was considered as a theta × HD cell. Among these theta × HD cells, they further identified theta skipping cells with theta skipping (TS) index *T S* > 0.1 (see below). These operations resulted in a total of 208 HD cells, among which 76 are normal HD cells without theta modulation, 77 are theta × HD cells and 30 of them are theta skipping cells. This dataset is publicly available at https://figshare.com/articles/dataset/_strong_Data_code_for_Lomi_et_al_2023_strong_/22802861.

### Decoding theta sweeps in the grid cell dataset with a state-space decoder

We adopted a state-space model [51] to decode theta sweeps in the grid cell dataset. Decoding of location was most reliable from Rat 1 (163 grid cells from two modules) and Rat 2 (544 grid cells from three modules) since cells are from multiple grid modules. The encoding part of the state-space model takes two inputs: the spike array of all spike-sorted neurons and the time-aligned 2D position data. The position data was binned into 4 × 4 cm spatial bins. We built the encoding model using a 2-ms temporal bin by up-sampling the position data from 100 Hz to 200 Hz. Only periods when the running speed exceeding 4 cm/s were considered in constructing the encoding model. Since theta sweeps closely track the animals’ physical locations, we defined latent movement dynamics as a random walk process (without considering uniform and fragmented dynamics as in their replay analysis [51]). The movement variance for the random walk process was computed as 10 times of the movement variance estimated from the whole position trajectory during recording. This amplification helps to increase the variance of the latent movement dynamics and makes the theta sweeps more obvious (setting the amplification value to 5 or 20 did not visually change the decoding results too much). For decoding, we used 5-fold cross-validation. In this process, we built the encoding model on 4 folds of the data and then decoded the sequences on the remaining fifth fold. This ensures that the data used for constructing a given encoding model were not used for decoding the representation. We repeated this for each fifth of the data.

### Sweep alternation score

To calculate the sweep alternation score, a three-sweep sliding window was used as in Vollan et al. [13]. Specifically, at the *i*^*th*^ time step, a triplet of sweep directions *α*_*i*−1:*i*+1_ was selected. Then the two angles between consecutive sweep pairs were calculated *a*_*i*_ = *α*_*i*_ − *α*_*i*−1_ and *b*_*i*_ = *α*_*i*+1_ − *α*_*i*_, and the alternation score at *i*^*th*^ time step was computed as:

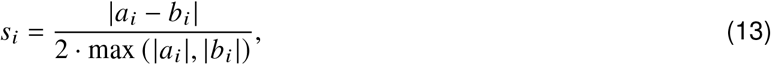

where | · | denotes the absolute value. The overall alternation score was finally calculated as the average of alternation scores across all simulation time bins: 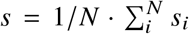, with *N* the total number of time bins. The alternation score is bound between 0 and 1, with higher values indicating more regular theta sweeps of left-right-left-right-left…

### Theta skipping index

To calculate the theta skipping index for each cell, we followed the method described in Brandon et al. [29]. First, we computed the spike time autocorrelogram (time range ±500 ms, bin width 5 ms). The autocorrelogram was further normalized by the peak value between 50 ms and 250 ms, and values above 1 (usually those around zero lag) was then clipped to 1, to ensure all the values are in a range of [0,1]. Second, the resulting autocorrelogram was fit with a cosine wave with a second interfering oscillation:

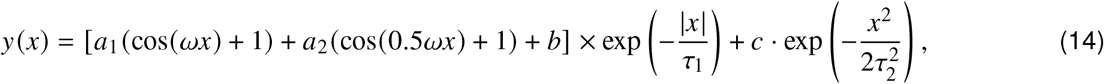

where *x* is the autocorrelation lag, *a*_1_ = [0, 1], *a*_2_ = [0, 1], *b* = [0, 1], *c* = [−1, 1], *ω* = [10*π*, 18*π*], *τ*_1_ = [0, 5] and *τ*_2_ = [0, 0.005] are the searching range for parameter fitting. Third, the theta skipping index was calculated as the difference between the first and second peaks in the autocorrelogram, normalized by the larger of the two:

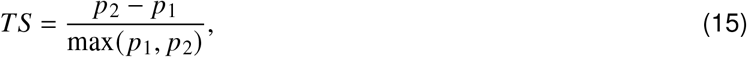

where *p*_1_ is the model value at one cycle with *x* = 2*π*/*ω* and *p*_2_ is the model value at two cycles with *x* = 4*π*/*ω*. This skipping index is bound between -1 and 1, with higher values indicating more theta skipping.

### Theta phase precession

To quantify theta phase precession in head direction (HD) cells recorded by Lomi et al. [18], we plotted the spike phase-head direction relationship for each cell. Spike phases were computed by applying the Hilbert transform to the bandpass-filtered local field potential (LFP) signal in the theta frequency range (6-12 Hz). Head directions were realigned to the preferred direction of the cell at the center of the x-axis (Fig. 7a). This realignment reduced the circular effect of the head direction signal on the x-axis and allowed us to perform circular-linear correlation [52] on the scatter plot, instead of circular-circular correlation.

Importantly, only periods of continuous head rotation were selected for constructing the scatter plot and conducting the circular-linear correlation analysis. Specifically, we required a minimum angular speed of 0.5 radians/second and a duration of at least 0.5 seconds. This results in approximately 15 degrees of continuous head rotation. In phase precession of place cells in a linear track environment, cells often develop directional tuning. Therefore, only uni-directional running periods are considered when analyzing the phase precession effect. However, HD cells fire during both clockwise and counterclockwise turns within their tuning field. To account for differences in phase precession due to tuning direction, we flipped the head directions during counterclockwise turns by subtracting the angle from 2*π*. This adjustment ensures that small head direction values on the x-axis consistently represent entry phases into the tuning field, and large values represent exit phases from the tuning field. Finally, we performed circular-linear correlation for each cell [52] with a restriction on the slope in the range of [-1,0] on the phase-direction scatter plots, and considered *p* < 0.05 as significant theta phase precession.

## Supplementary figures

**Figure S1:**
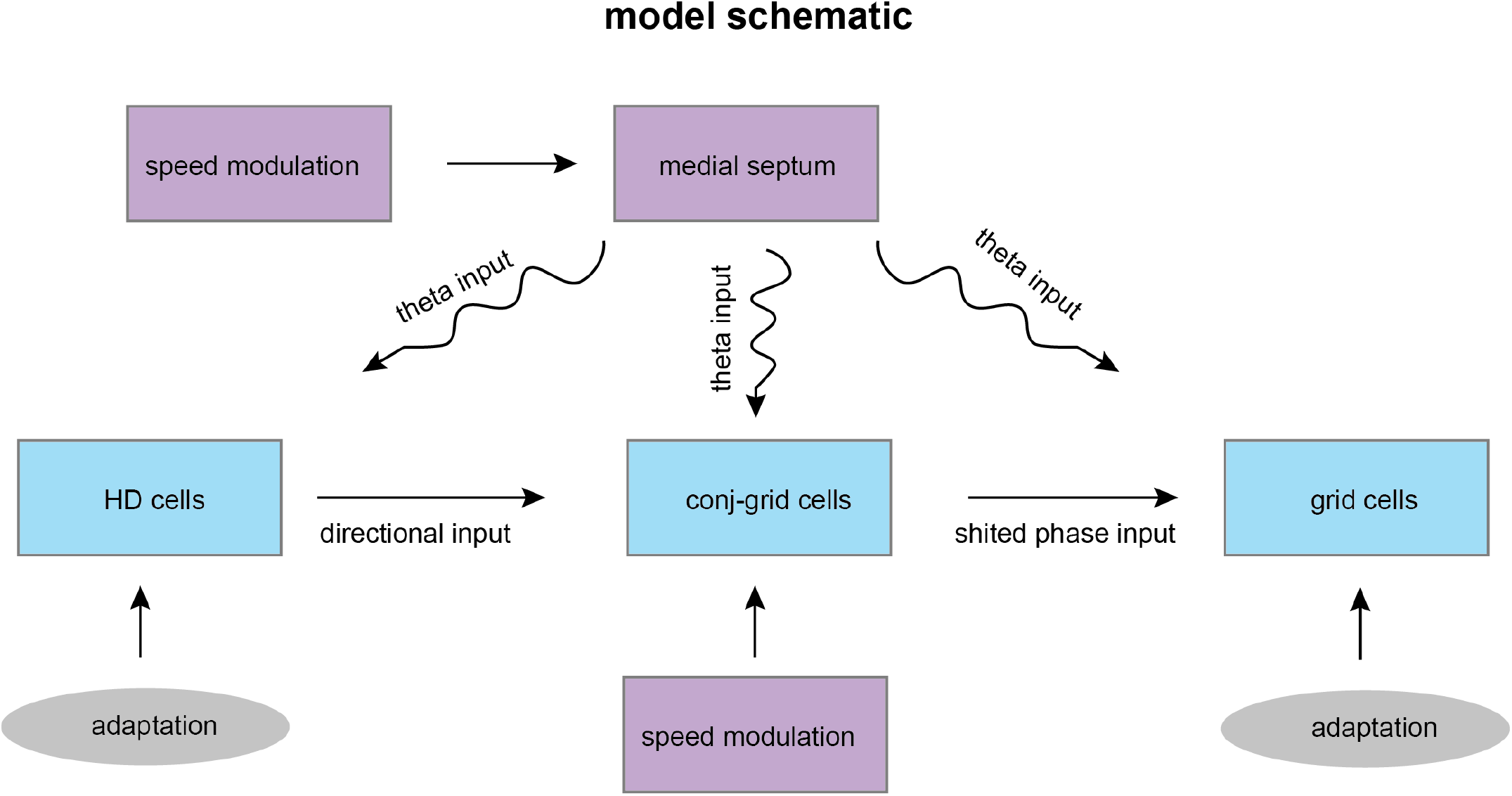
The schematic of the computational model. Blue blocks: simulated neurons including head direction, conjunctive direction x grid and pure grid cells. Purple blocks: speed modulation of theta oscillation from medial septum. Grey blocks: firing rate adaptation.

**Figure S2:**
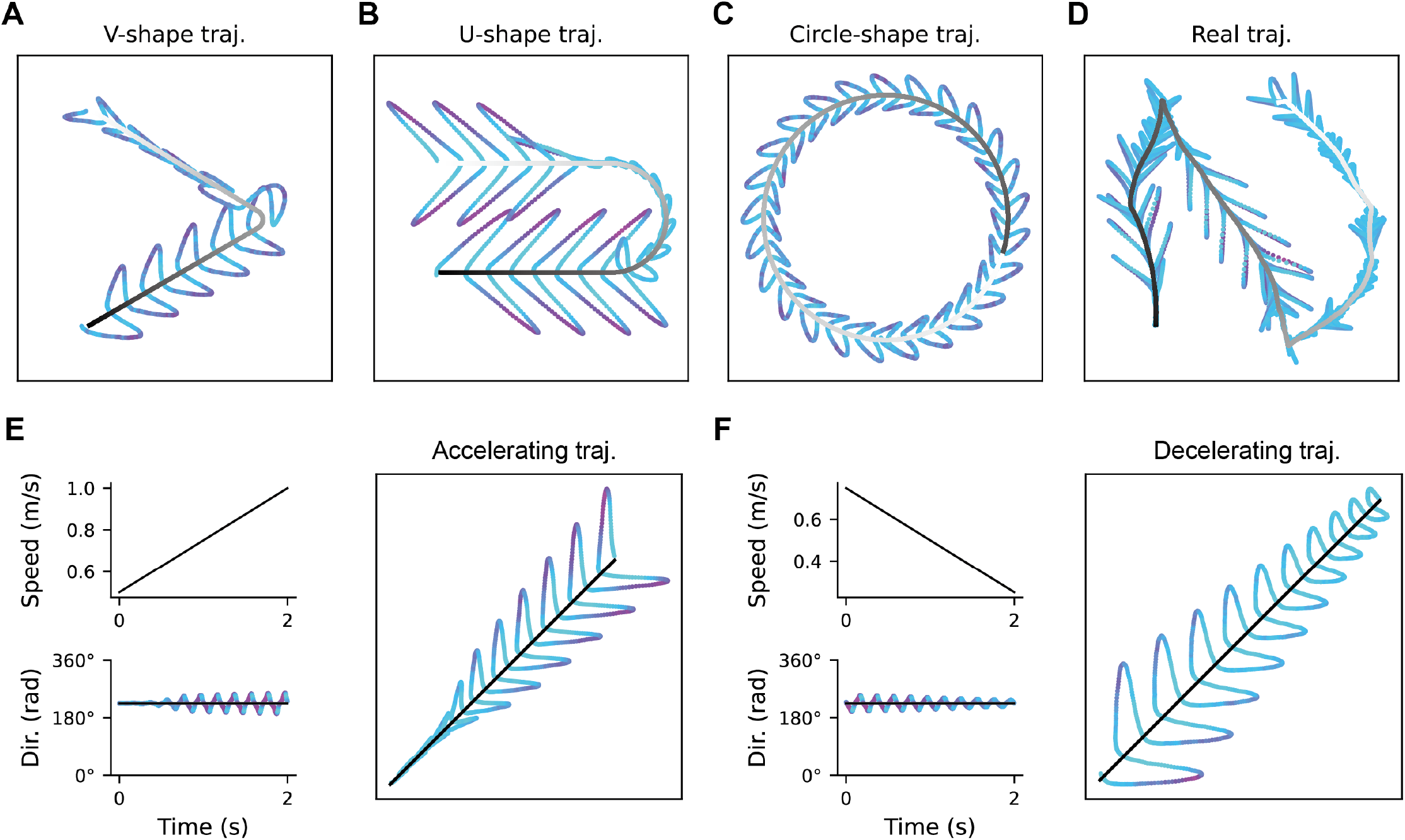
Theta sweeps when animal runs along non-stereotypical trajectories. **a**. Theta sweeps on a V-shape trajectory. **b**. Theta sweeps on a U-shape trajectory. **c**. Theta sweeps on a round-shape trajectory. **d**. Theta sweeps on a trajectory simulated with a random walk process (also shown in Fig. 3c). **e**. Theta sweeps on a straight trajectory when the animal’s speed is accelerating. Left top: the animal’s speed as a function of time. Left bottom: increased angle of the internal direction sweeps over an accelerating time period. Right: increased length and angle of the location sweeps over an accelerating time period. **f**. Decreased angle of the internal direction sweeps and decreased length and angle of the internal location sweeps over a decelerating time period.

**Figure S3:**
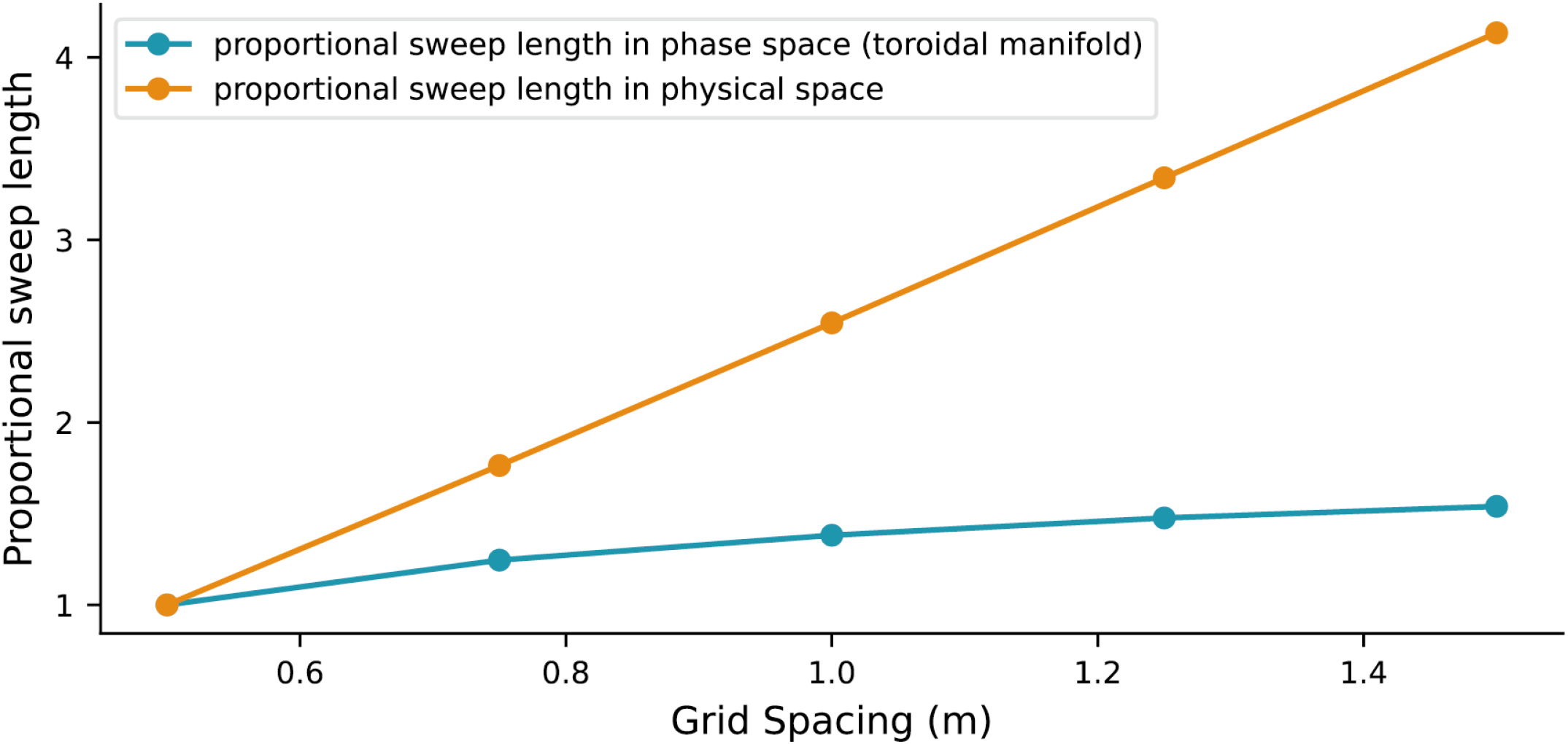
Theta sweep length on the toroidal manifold and in the physical space as a function of grid spacing. Since the unit of measurement in the phase space and the physical space are different, the sweep length was normalized by the sweep length with the one simulated with the smallest grid spacing.

**Figure S4:**
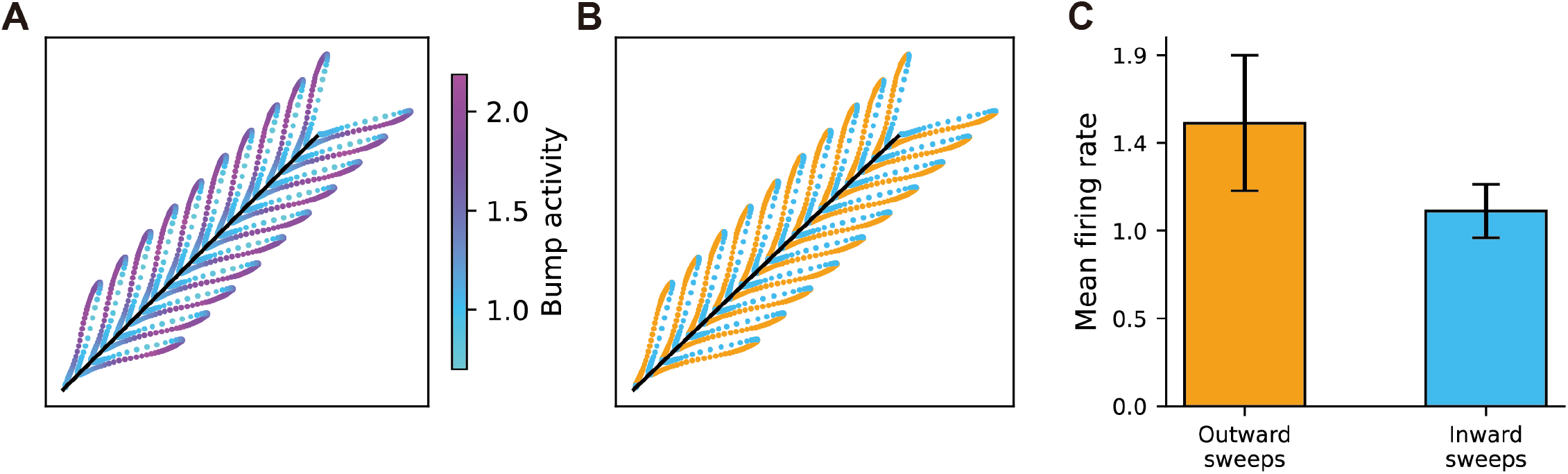
Mean firing rate of grid cells during outward sweeps and inward sweeps. **a**. Left-right theta sweeps in the grid cells. With more purple dots represent higher firing rates of the network bump activity. **b**. Left-right theta sweeps during outward phase (orange) and inward phase (blue). **c**. The averaged network bump activity during outward phase (orange) and inward phase (blue).

**Figure S5:**
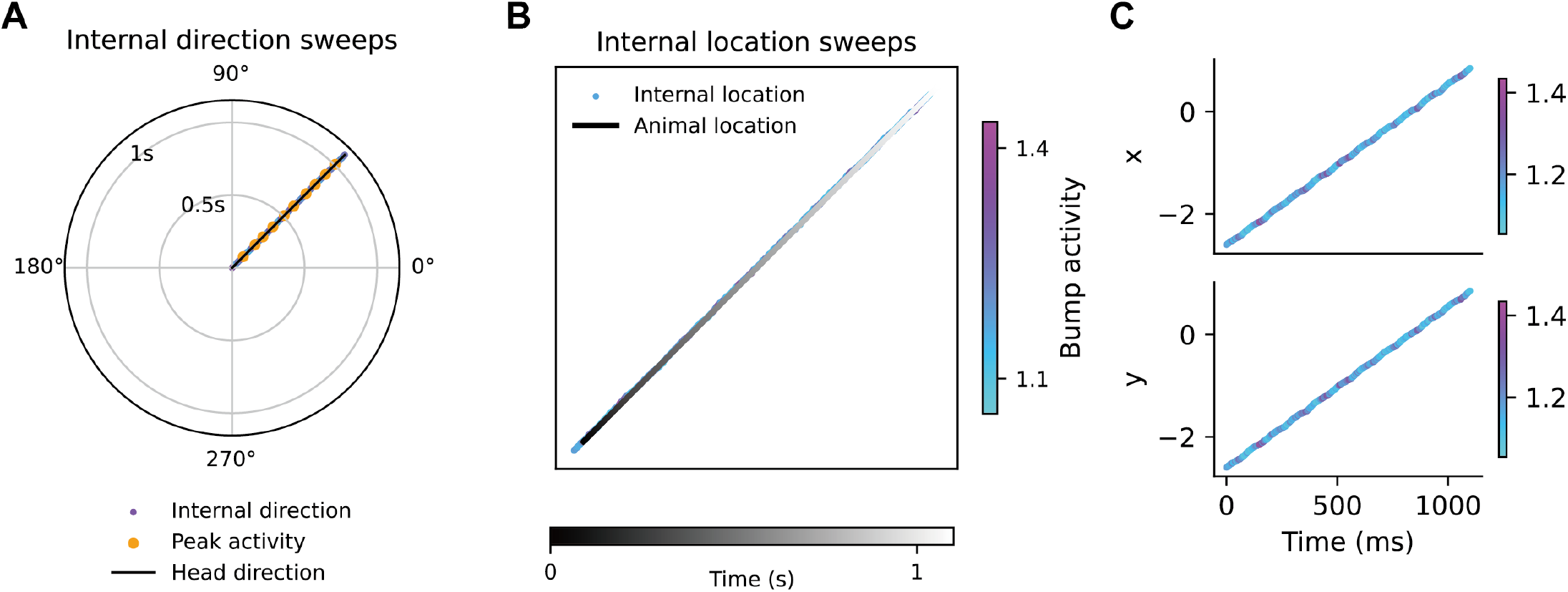
Blocking firing rate adaptation eliminates theta sweeps in both head direction cells and grid cells. **a**. Without firing rate adaptation, the internal direction in the HD attractor tracks the animal’s head direction. **b**. Without firing rate adaptation, the internal location in the GC attractor also tracks the animal’s physical location. **c**. Separate plots of the two dimensions of the internal location in **b** over time.

**Figure S6:**
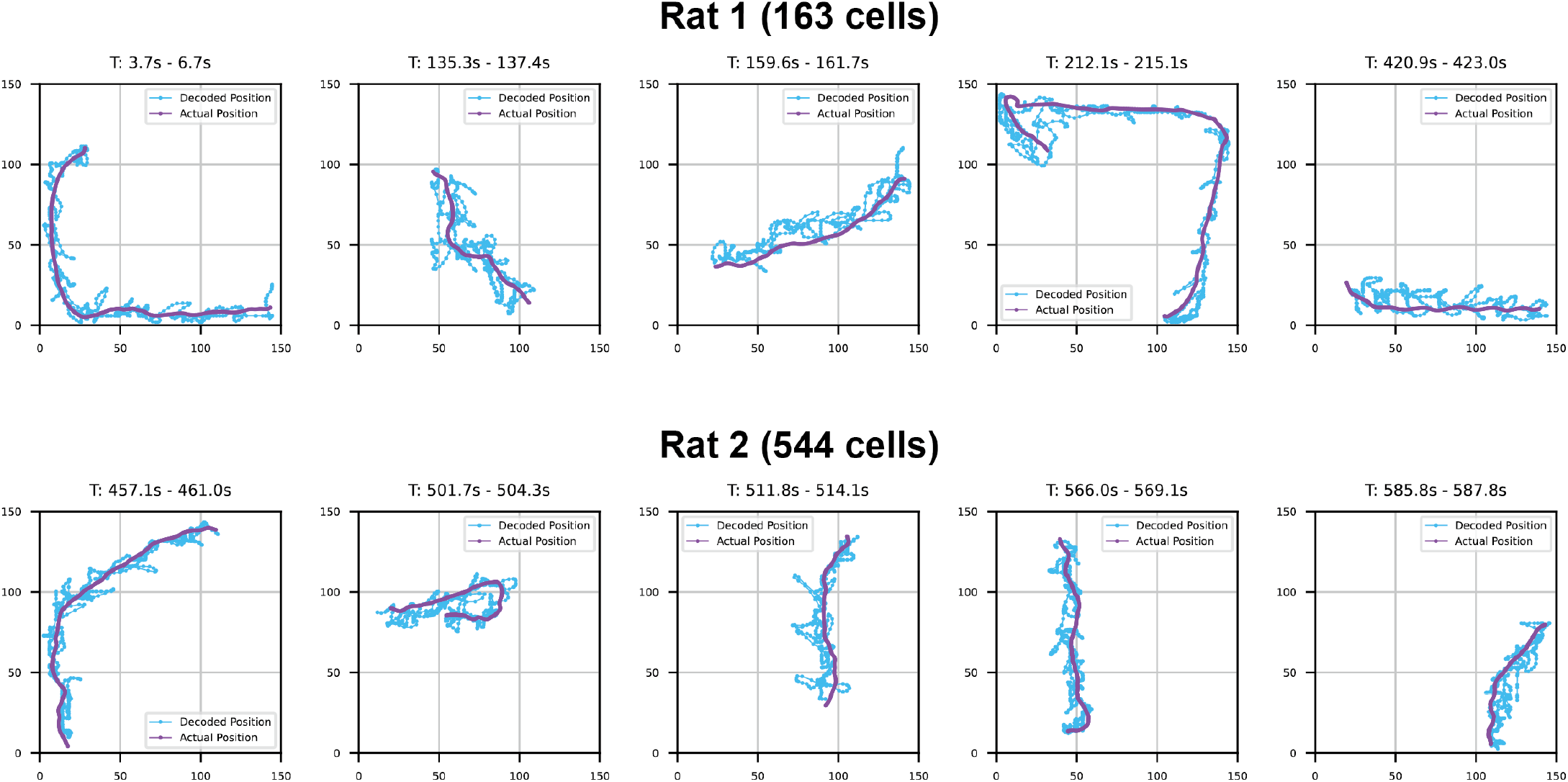
Theta sweeps decoded with a state-space model [51] from the dataset provided by Gardner et al. [24]. Also see Video.3. Rat 1 has two grid modules with 164 cells in total. Rat 2 has three grid modules with 544 cells in total. For each rat, we randomly selected 5 fast run periods to decode theta sweeps. Note that the state-space decoder was 5-fold cross-validated, meaning that data for encoding part was not seen by data for decoding.

## Supplementary tables

**Table 1:**
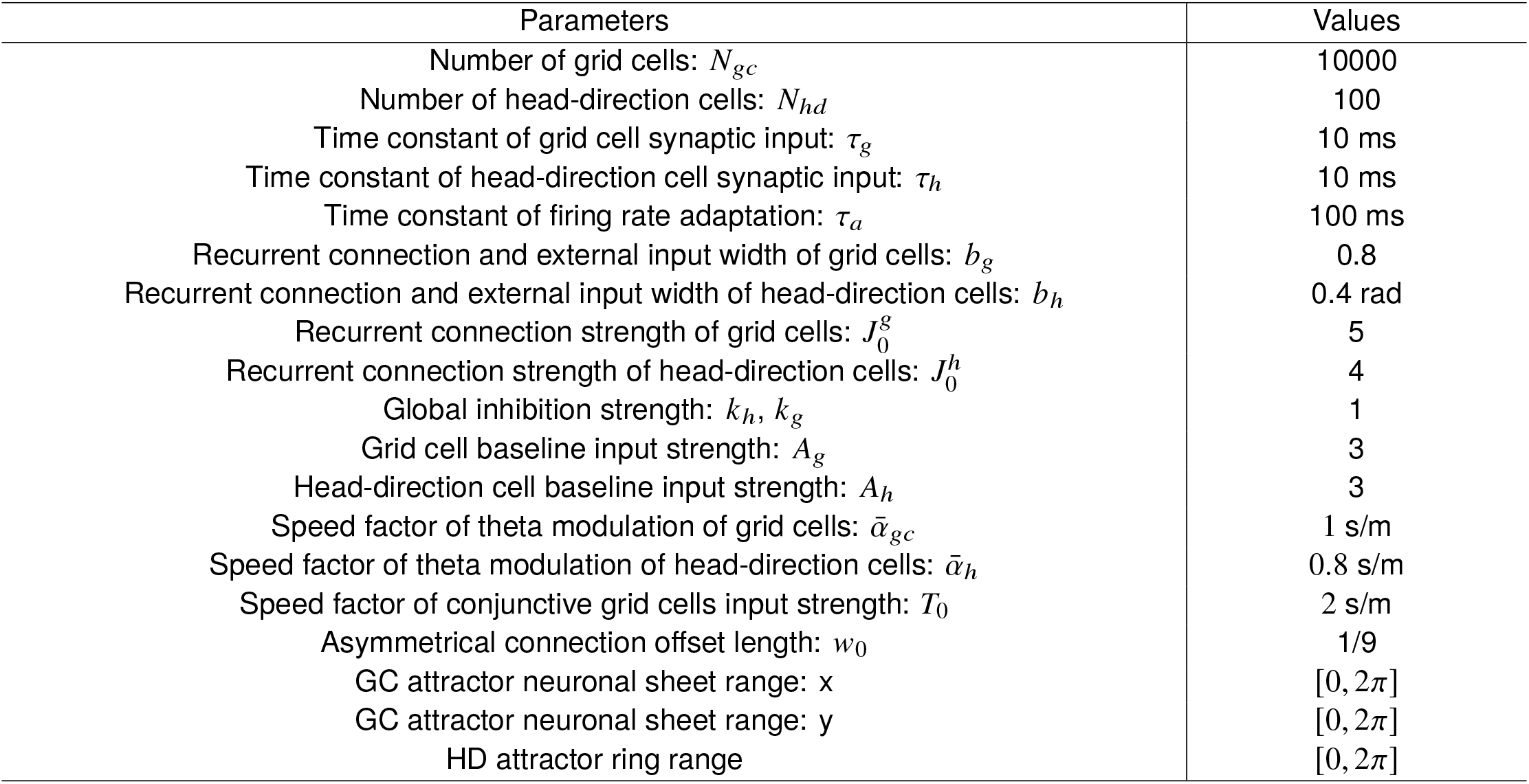
Commonly used parameters across simulations. Note that these parameters were not tuned in effort. A large range of parameter values can lead to similar phenomena showed in this paper.

| Parameters | Values |
| --- | --- |
| Number of grid cells: $N_{gc}$ | 10000 |
| Number of head-direction cells: $N_{hd}$ | 100 |
| Time constant of grid cell synaptic input: $\tau_g$ | 10 ms |
| Time constant of head-direction cell synaptic input: $\tau_h$ | 10 ms |
| Time constant of firing rate adaptation: $\tau_a$ | 100 ms |
| Recurrent connection and external input width of grid cells: $b_g$ | 0.8 |
| Recurrent connection and external input width of head-direction cells: $b_h$ | 0.4 rad |
| Recurrent connection strength of grid cells: $J_0^g$ | 5 |
| Recurrent connection strength of head-direction cells: $J_0^h$ | 4 |
| Global inhibition strength: $k_h, k_g$ | 1 |
| Grid cell baseline input strength: $A_g$ | 3 |
| Head-direction cell baseline input strength: $A_h$ | 3 |
| Speed factor of theta modulation of grid cells: $\bar{\alpha}_{gc}$ | 1 s/m |
| Speed factor of theta modulation of head-direction cells: $\bar{\alpha}_h$ | 0.8 s/m |
| Speed factor of conjunctive grid cells input strength: $T_0$ | 2 s/m |
| Asymmetrical connection offset length: $w_0$ | 1/9 |
| GC attractor neuronal sheet range: $x$ | $[0, 2\pi]$ |
| GC attractor neuronal sheet range: $y$ | $[0, 2\pi]$ |
| HD attractor ring range | $[0, 2\pi]$ |

**Table 2:** Figure specific parameter values for moving speed *v*, grid-cell adaptation strength *m*_*g*_, HD-cell adaptation strength *m*_*h*_, and grid spacing *λ*.

| Figures/parameters | $v$ | $m_g$ | $m_h$ | $\lambda$ |
| --- | --- | --- | --- | --- |
| Figure 2 & 3a-3f | 1 m/s | 1. | 1.2 | 1 m |
| Figure 3g-3h | 0.5 m/s | 1.5 | 1.2 | 0.3 : 0.1 : 1.5 m |
| Figure 4 | 0.1 : 0.225 : 1 m/s | 1 | 1.2 | 1 m |
| Figure 5 | 0.5 m/s | 0.7:0.09:1.8 | 0.7:0.09:1.8 | 1 m |

## Supplementary videos

Video 1: Theta sweeps in the model with a real running trajectory consisting of straight runs, turns, immobile periods.

Video 2: Theta sweeps in the model across three grid modules.

Video 3: Decoded theta sweeps from real data collected by Gardner et al. [24].

